# Identification of a novel family of benzimidazole species-selective Complex I inhibitors as potential anthelmintics

**DOI:** 10.1101/2022.09.12.507634

**Authors:** Taylor Davie, Xènia Serrat, Jamie Snider, Igor Štagljar, Hiroyuki Hirano, Nobumoto Watanabe, Hiroyuki Osada, Andrew G Fraser

## Abstract

Soil-transmitted helminths (STHs) including *Ascaris*, hookworm, and whipworm are major human pathogens infecting over a billion people worldwide^1,2^. There are few existing classes of anthelmintics and resistance is increasing^3–5^ — there is thus an urgent need for new classes of these drugs. Here we focus on identifying compounds that interfere with the unusual anaerobic metabolism that STHs use to survive the highly hypoxic conditions of the host gut^6–9^. This requires rhodoquinone (RQ), a quinone electron carrier that is not made or used by the STH hosts^10^. We previously showed that *C. elegans* also uses this rhodoquinone-dependent metabolism (RQDM)^11^ and established a high throughput assay for RQDM^11^. We screened a collection of 480 natural products for compounds that kill worms specifically when they rely on RQDM — these 480 are representatives of a full library of ~25,000 natural products and derivatives^12,13^. We identify several classes of compound including a novel family of species selective inhibitors of Complex I. These Complex I inhibitors are based on a benzimidazole core but unlike commercial benzimidazole anthelmintics they do not target microtubules^14–17^. We screened over 1,200 benzimidazoles and identify the key structural requirements for species selective Complex I inhibition. We suggest that these novel benzimidazole species-selective Complex I inhibitors may be potential anthelmintics.

## Intro

Soil-transmitted helminths (STHs from here on) are major pathogens of humans and livestock^1,2^. Over a billion humans are infected by STHs including roundworm (*Ascaris lumbricoides*, hookworm (*Necator americanus* and *Anclyostoma duodenale*), and whipworm (*Trichuris trichiura*). These infections result in malnutrition, malaise and weakness, and can cause developmental defects and impaired growth in children^18^. In addition, STHs infect a high proportion of livestock leading to reduced yield. This is a particular problem in poorer communities where such losses can have major health and economic consequences. There are excellent frontline anthelmintics including benzimidazoles (e.g. albendazole and mebendazole)^19,20^ and macrocyclic lactones (e.g. ivermectin)^21,22^. However, there are few classes of commercial anthelmintics and resistance to these drugs is widespread in livestock and is now occurring in human parasites as well. There is thus an urgent need for new classes of anthelmintic drugs to control and treat these major pathogens which present a key challenge in global health.

Any effective anthelmintic must target the STH without harming the vertebrate host. One way to do this is to develop drugs that target a process that is essential for the STH but absent from the host and that is the approach we take here. We focus on a unique aspect of STH metabolism — rhodoquinone-dependent metabolism (RQDM)^6,7,9,23^. During the stages of the lifecycle when the STH lives in the soil outside the host, it generates energy using oxidative phosphorylation. Electrons enter the mitochondrial electron transport chain (ETC) either at Complex I or via quinone-coupled dehydrogenases (QDHs) like Complex II, succinate dehydrogenase. They are transferred to ubiquinone (UQ) and then pass through Complex III and Complex IV where they ultimately are accepted by molecular oxygen as the terminal electron acceptor. As the electrons flow through the ETC, protons are translocated across the mitochondrial membrane, establishing a proton motive force which powers ATP synthesis by the F0F1-ATP synthase, Complex V (Fig 1a). This UQ-using oxidative ETC is identical between host and parasite. However, many STHs must survive extended periods in the highly anaerobic environment of the host gut — adult *Ascaris*, for example lives for months in these conditions. Without available oxygen as the terminal electron acceptor, the STHs cannot use the aerobic UQ-coupled ETC. However, STHs have a key adaptation that still allows them to use a rewired form of the ETC and it is this that provides the potential target for anthelmintics. Electrons still enter the ETC from NADH into Complex I and onto the quinone pool. Rather than flow through to Complex IV, they exit the ETC at Complex II which now acts as a fumarate reductase rather than a succinate dehydrogenase (Fig 1b). This allows fumarate to be used as a terminal electron acceptor. Crucially, UQ cannot be used as an electron carrier to power the fumarate reductase activity — instead STHs use rhodoquinone (RQ), a highly related quinone^24–26^. Since only STHs make and use RQ, RQ synthesis and RQDM provide a critical target that differs between host and parasite. If we could identify drugs that specifically target RQ synthesis or RQDM, they should kill the parasite without affecting the host — this is our goal here.

**Figure 1:**
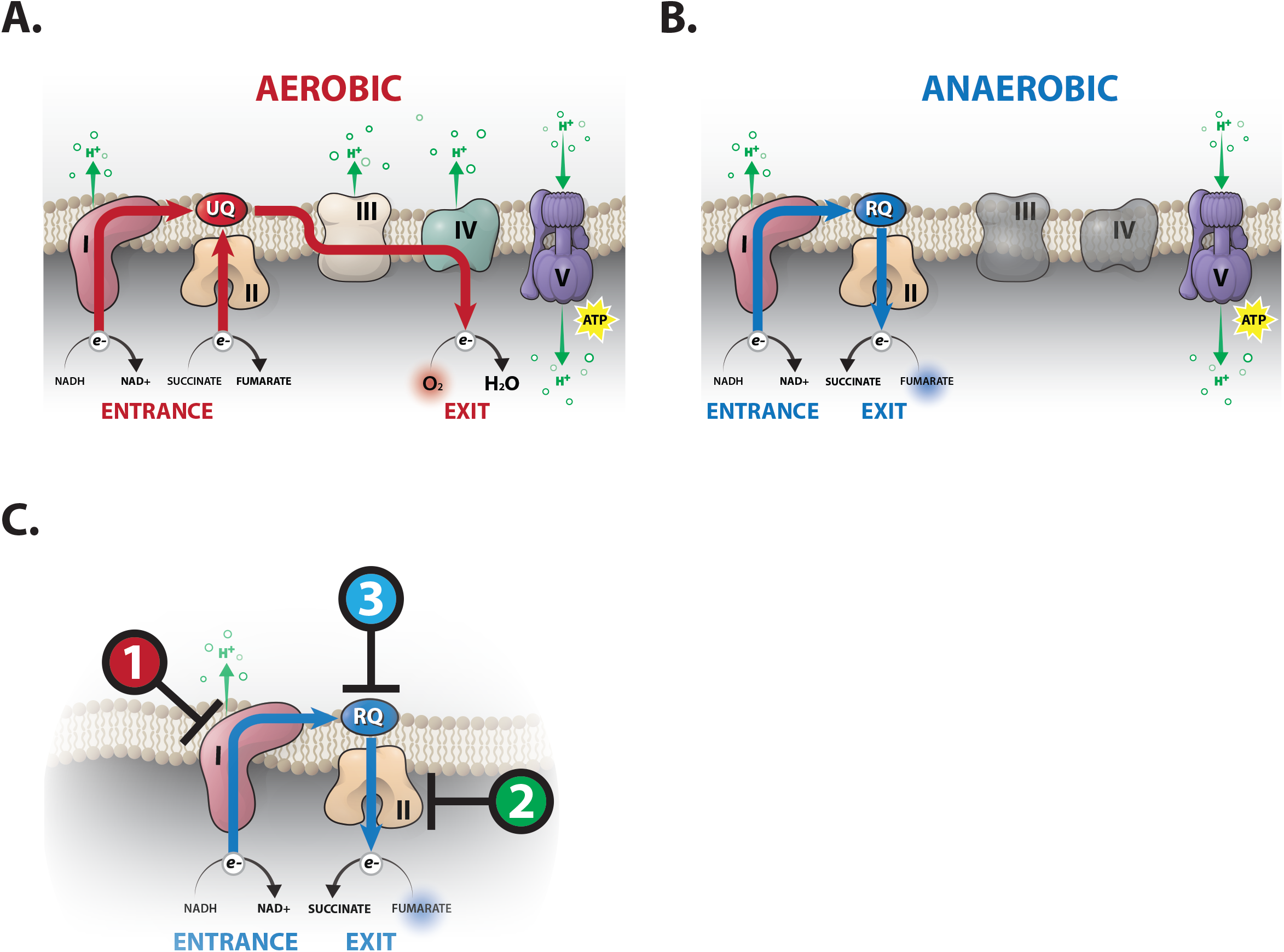
Ubiquinone-coupled aerobic and rhodoquinone-coupled anaerobic electron transport chain (ETC). **A. Ubiquinone-coupled aerobic ETC**. Electrons enter the quinone pool via Complex I or Complex II and exit via Complex IV — oxygen is the terminal electron acceptor. Ubiquinone is the quinone electron carrier and protons are pumped across the mitochondrial membrane by Complex I, III and IV; ATP is synthesised by Complex V powered by the proton motive force. **B. Rhodoquinone-coupled anaerobic ETC.** Electrons enter the quinone pool via Complex I only and exit via Complex II which acts as a fumarate reductase — fumarate is the terminal electron acceptor. Rhodoquinone is the quinone electron carrier and only Complex I pumps protons. **C. Key targets for inhibitors of Rhodoquinone-coupled anaerobic ETC.** Compounds that target Complex I (sole electron entry point, sole proton pump), Complex II (key electron exit point), and RQ synthesis (key quinone electron carrier) are likely to act as anthelmintics.

We previously showed that the free-living nematode *C. elegans* makes and uses RQ^11^. We established a simple assay which allows us to measure RQDM *in vivo* and used this to dissect the pathway for RQ synthesis^11^ and the key molecular switch that determines whether nematodes make UQ or RQ^27^. In brief, tryptophan is metabolised via the kynurenine pathway to generate 3-hydroxyanthranilate (3HA). 3HA is then used as a substrate by the polyprenyl transferase COQ-2 in the critical commitment step in RQ synthesis^11,28^. Mutations in the kynurenine pathway such as a *kynu-1* null mutation block 3HA generation and hence block RQ synthesis. Wild-type worms, in which ~10% of the quinone pool is RQ^11,28^, can readily survive treatment for 15hrs with 200μM potassium cyanide (KCN) but *kynu-1* worms that entirely lack RQ cannot survive this. We further showed that inhibitors of Complex I or Complex II, which prevent either electron entry or exit from the RQ-dependent ETC, or mutations in the quinone-binding pocket of Complex II that prevent RQ binding kill worms in the presence of 200μM KCN (Fig 1c)^11^. *C. elegans* is thus an excellent model for RQDM and it is relatively easy to carry out high throughput drug screens *in vivo* in *C. elegans.*

In this study, we screen a collection of 480 natural products and their derivatives^12,13^ for compounds that affect our *C. elegans* model of RQDM. We identified multiple distinct structural groups of potential RQDM inhibitors and showed that several of these have highly specific effects on individual ETC complexes. These include a species-selective Complex II inhibitor, siccanin, that had not been previously shown to be active in nematodes, and a family of structurally related benzimidazole compounds that specifically and potently target Complex I. We showed Complex I inhibition is a novel activity for benzimidazoles — no commercial benzimidazoles have any effect on Complex I activity. Furthermore, we identified a small subset that show good species selectivity, inhibiting *C. elegans* Complex I with >10-fold lower doses than bovine or murine Complex I and that have no detectable effect on growth in normoxia. We have thus identified a new family of species-selective Complex I inhibitors that potently kill *C. elegans* under conditions where they require RQDM for survival. These are potential anthelmintics that may similarly kill STHs in the host gut where they rely on RQDM for survival.

## Results

### A screen in *C. elegans* for natural products that inhibit RQ-dependent metabolism

*C. elegans* is highly related to STHs^29^ and *C. elegans* makes and uses RQ^11,28^. It is thus an excellent system for screens to identify compounds that specifically block RQ synthesis and RQ-dependent metabolism (RQDM). We previously showed that *C. elegans* requires RQ synthesis and RQDM to survive exposure to KCN for 15hrs^11^. This observation is the basis for a previously published image-based movement assay for RQDM. In outline, we treat *C. elegans* L1 larvae with 200 μM KCN for 15hrs — wild-type worms that can produce RQ are immobile but alive and if we remove the KCN by dilution they rapidly recover movement (Fig 2a). However, worms that cannot make RQ (e.g. that have mutations in the kynurenine pathway (Fig 2a)) or that cannot carry out RQDM (e.g. due to inhibition of Complex I or Complex II (Fig 2b)) are dead after 15hr KCN treatment and hence no recovery of movement is seen. This allows a simple drug screen for products that specifically kill *C. elegans* when they require RQDM for survival. Worms are treated with 200 μM KCN with either test compounds or DMSO only controls and KCN is removed after 15hrs and movement measured 3hrs later. If worms recover movement similar to controls, then the compound does not affect RQDM. However, if worms are dead after 15hr KCN treatment in the presence of the test compound, no recovery is seen — that compound is thus a potential inhibitor of RQDM. Finally, to ensure that the compound’s effect is specific for RQDM and not some more general effect on worm viability, we also examine the effect of each compound on the growth and viability of worms in normoxia where they do not require RQ and do not use RQDM. If a compound has no effect on growth and viability in normoxia but is strongly lethal in our RQDM assay, this is a potential specific inhibitor of RQ synthesis or RQDM. Our goal here is to identify such compounds and characterise their effects on RQDM — they are candidates as anthelmintics that act via inhibition of the anaerobic RQDM that allows STHs to survive in the host gut.

**Figure 2:**
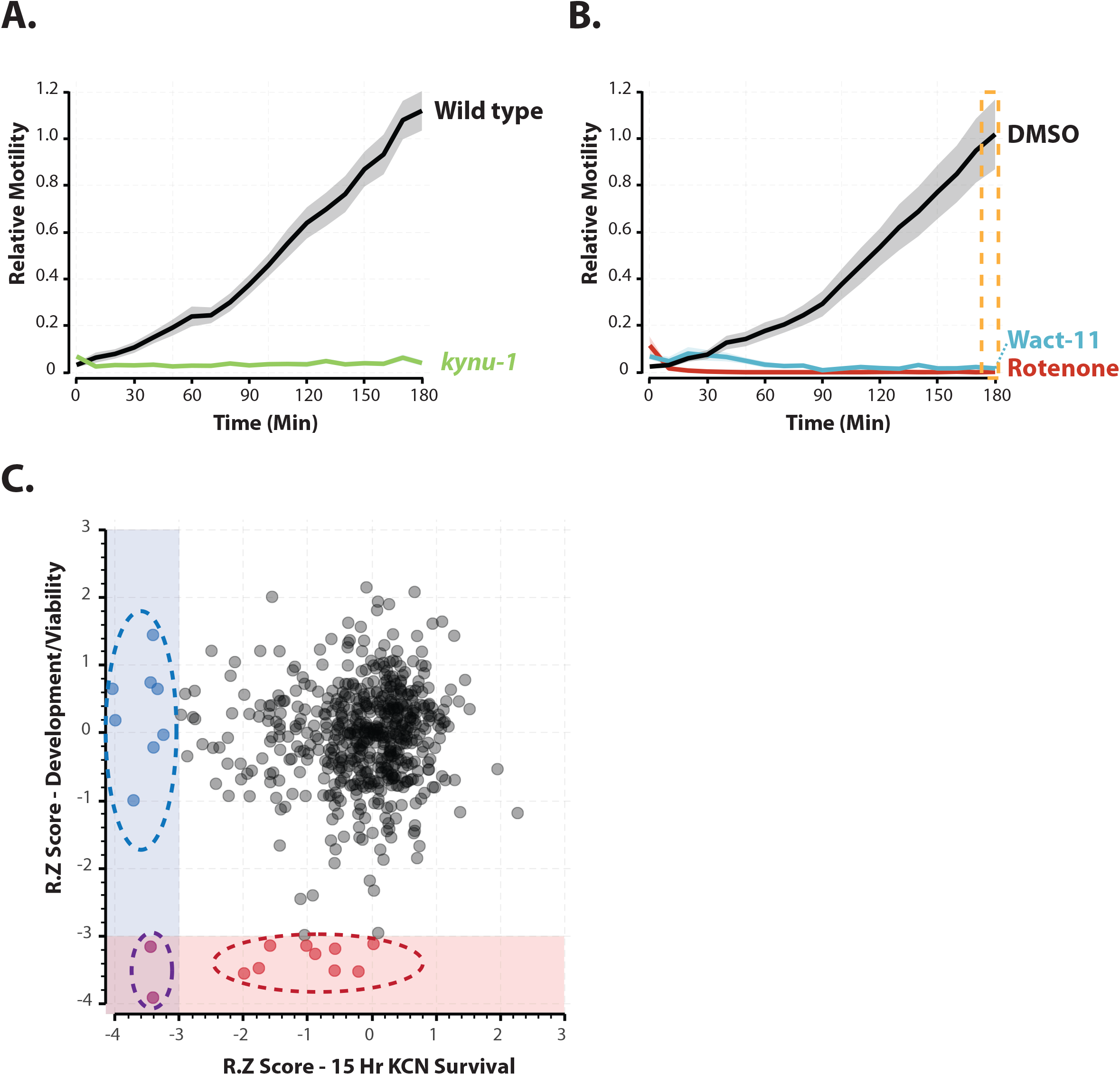
Screens to identify inhibitors of RQ-dependent metabolism (RQDM). **A. Phenotype of *C. elegans* lacking RQDM.** Wild-type worms exposed to 200 μM KCN for 15hrs are immobile but rapidly recover full movement when KCN is removed (black curve). *kynu-1* null mutants that lack RQ and thus cannot carry out RQDM are dead after 15 hr KCN exposure and thus cannot recover movement. The ability to recover movement after 15 hrs of 200 μM KCN exposure is thus a simple assay for the ability to carry out RQDM. Data are the mean of three biological replicates; errors are S.E.M. **B. Effect of ETC inhibitors on RQDM.** Inhibition of Complex I (rotenone) or Complex II (wact-11) prevents worms carrying out RQDM. **C. Data from screens of ~480 natural products (NPDs) in RQDM assays and growth assays.** Worms were treated with each NP at 50 μM and screened in 2 assays — the RQDM assay and a quantitative growth assay (see Materials and Methods). Each dot represents a single NP and is the mean of 3 biological replicates. Data are presented as modified z-scores relative to the median. 8 NPs showed significant effects (r.z<-3) in the RQDM assay alone; 9 in the growth assay alone; and 2 in both assays.

We screened a library of 480 natural products and their derivatives from the Riken Natural Product Depository (RIKEN NPDepo)^12,13^ in our KCN-based assay for RQDM and also in a more traditional *C. elegans* normoxic growth and viability assay. 80 of these natural products (authentic library) were known molecules with previously characterized biological activities. The remaining 400 compounds (pilot library) are a subset of a larger library of 25,000 diverse natural product derivatives (NPDs) — each compound represents a family of structurally related NPDs. This allows a rapid initial primary screen covering the broad structural space of the entire NP collection and, once hits are identified, structurally related NPDs are then screened in secondary assays. This both helps define the structural requirements for the biological effects of the primary hits as well as identifying any related compounds with greater potency. The overall results are shown in Fig2c. For both assays, each compound was screened in triplicate at 50 *μ*M and the mean value was expressed as a modified z-score. Compounds with modified z-scores < −3 (i.e. < 3X median absolution deviation (MAD)) were identified as hits; all data are shown in Supp Table 1.

We identified 9 compounds that affect normoxic growth. Five of these were natural products with known activities and included nocodazole, an inhibitor of microtubule polymerisation^30^; staurosporine, a broad-spectrum kinase inhibitor^31^; camptothecin, a DNA topoisomerase inhibitor^32^; reveromycin A, an isoleucyl-tRNA synthetase inhibitor^33^; and aristolochic acid, a known mutagen^34^. These drugs all target core essential cellular components and the observed lethality is consistent with this. We note these are all known to have toxic effects on mammalian cells and thus are not suitable as anthelmintics. In addition to the hits with known bioactivities, we identified four compounds with novel scaffolds that inhibited *C. elegans* growth and viability. Several of these compounds show no activity in HEK293 cells but readily kill *C. elegans.* Future work will involve screening related NPDs to further explore the utility of these compounds as potential anthelmintics. For the purposes of this study, we focused on the 8 compounds that showed little effect on growth in normoxia but killed worms specifically when they were relying on RQDM for survival. These are candidate RQDM inhibitors and we refer to these as ‘RQDM hits’.

### NPD screens reveal siccanin as a species-selective inhibitor of Succinate Dehydrogenase

The NP library contains two groups of compounds: the larger group (pilot library) is of NPDs with no previous reported bioactivity, and a second smaller group (authentic library) includes compounds that had previously been shown to be active in at least one previous screen. This latter group yielded two potent RQDM hits known to affect oxidative phosphorylation. Niclosamide is a salicylanilide anthelmintic used to treat tapeworm infections^35^. Niclosamide acts as an ionophore and exerts its lethal effects by collapsing mitochondrial membrane potential^36^. We had previously found that other salicylanilide anthelmintics such as closantel are powerful inhibitors of RQDM (data not shown) and thus this hit is expected.

The second hit, siccanin, is less well characterised but has been reported to target Complex II, succinate dehydrogenase, in a species selective manner^37,38^. Complex II is the principal exit point for electrons during RQDM^6,8^ and we had previously shown that inhibition of Complex II can kill *C. elegans* when they rely on RQDM for energy generation. The species selectivity of siccanin of Complex II inhibition had previously been shown in fungi, bacteria, and mammals^37^, but there are no reports of its effectiveness in any helminth or nematode. We find that siccanin inhibits *C. elegans* Complex II and that this is highly species selective (Fig 3A). The IC50 vs *C. elegans* Complex II is ~ 8 nM whereas IC50 vs mouse Complex II is ~18 *μ*M (Fig 3A) equating to >1000-fold difference. The potency of complex II inhibition by siccanin is equivalent to wact-11, a nematode-specific Complex II inhibitor structurally related to the anthelmintic fluopyram^39,40^.

**Figure 3:**
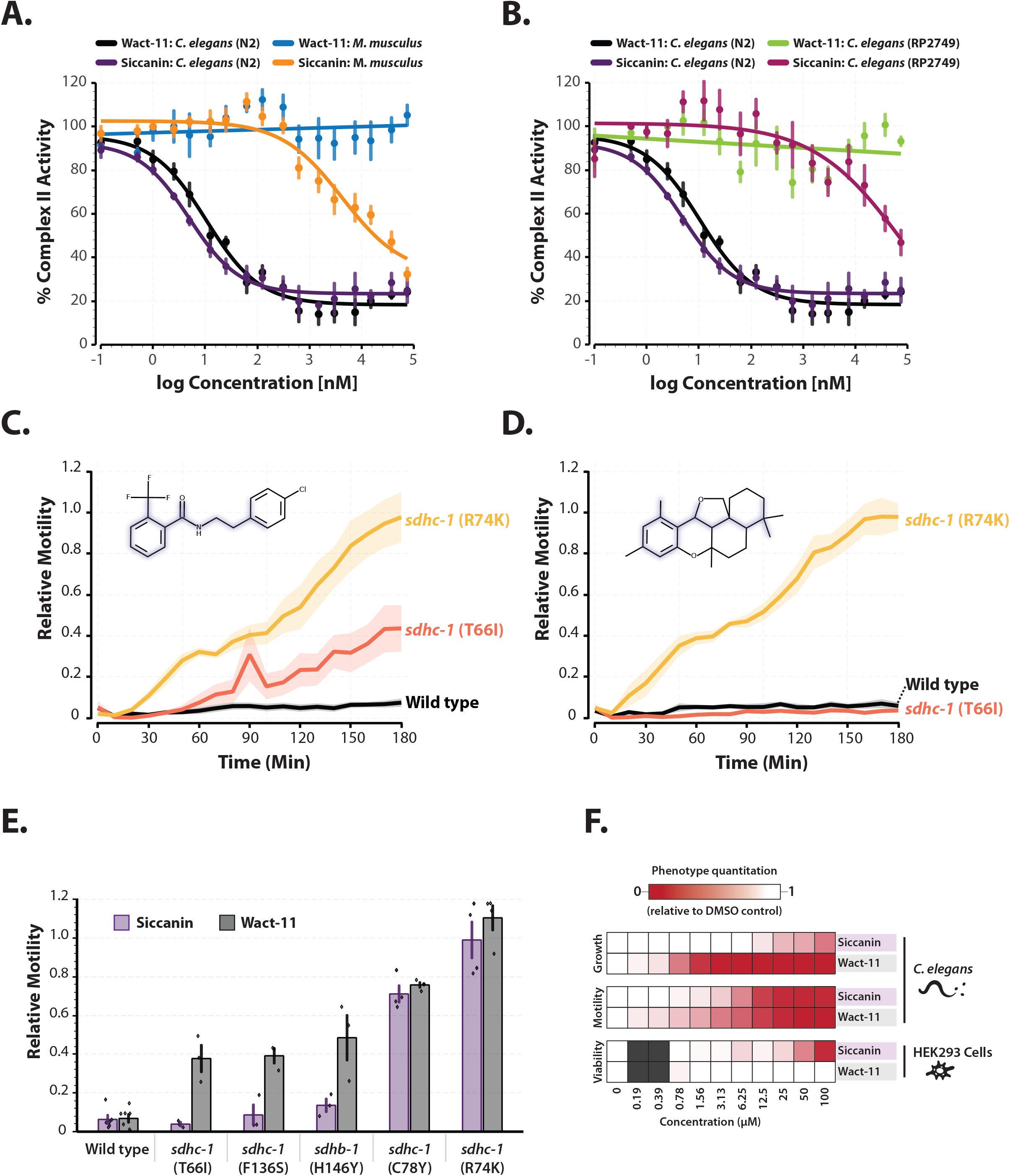
The antifungal Siccanin inhibits *C. elegans* Succinate Dehydrogenase (Complex II) **A.** Complex II inhibitory dose-response curves for siccanin and wact-11 against complex II from wild-type (N2) worm mitochondria (purple and black) and *M. musculus* liver mitochondria (yellow and blue). Assays were conducted as described in Materials and Methods; all data are the mean of 4 biological repeats and errors are the S.E.M. **B. Mutation of the quinone binding pocket of complex II causes insensitivity to Siccanin.** Complex II inhibitory dose-response curves for siccanin and wact-11 against complex II from *sdhc-1* (RP2749) mutant worm mitochondria (majenta and green). The RP2749 strain is homozygous for a mutation in the quinone binding pocket of Complex II and causes resistance to wact-11. **C and D. The mutational spectrum of siccanin and wact-11 resistance is distinct.** Two strains homozygous for mutations in the SDHC-1 subunit of Complex II (R47K in yellow and T66I in red) were treated with 200 μM KCN and either wact-11 (C) or siccanin (D) at 25 μM for 15hrs, after which KCN was diluted and worm movement measured for 3hrs. In all cases data are the mean of 3 biological replicates. **E.** Worm motility post 3 hours KCN dilution for wact-11 family resistant mutants that harbour different mutations in the ubiquinone binding pocket; different mutations show altered susceptibility to complex II inhibition by siccanin or wact-11. **F.** Activity profiles of siccanin and wact-11 against *C. elegans* (growth and motility) and HEK293 cells (viability). While wact-11 activity is specific to *C. elegans*, siccanin shows less selectivity.

To test whether siccanin binds to a similar site on Complex II as wact-11, we tested whether point mutations in the quinone binding pocket that are known to prevent wact-11 binding^39^, and thus cause resistance to wact-11, also affect siccanin inhibition. We found that the RP2749 strain that has a single mutation in the quinone binding pocket of Complex II is insensitive to both wact-11 and siccanin (Fig 3b), confirming the binding site and direct mode of action of siccanin (Fig 3B-D). We note that the binding is subtly different for wact-11 and siccanin — for example, a T66I mutation in *sdhc-1*, a subunit of Complex II, causes resistance to wact-11 but not to siccanin (Fig 3C and 3D) and other mutations show similar effects (Fig 3E). This suggests that while they share a target, some mutations that could cause resistance to fluopyram/wact-11 analogues might not affect siccanin killing. However, we find that siccanin has lower species selectivity than wact-11, lower potency in growth assays and higher toxicity in mammalian cells (Fig 3F). It is thus unlikely to be an improvement on previously identified Complex II inhibitors as an anthelmintic. Nonetheless we were encouraged that the 2 RQDM hits in the characterised NPD set had clear roles in electron transport and mitochondrial function and turned our attention to the previously uncharacterised NP products.

### NPD screens identify a structurally related group of Complex I inhibitors

The NPD screens identified 7 primary hits as NPDs with no previously described bioactivity that were lethal to *C. elegans* when they rely on RQDM for energy generation. We retested all available derivatives in the whole NP library^12,13^ of these 7 primary hits in the same RQDM assay and identified 7 clusters of structurally related hits (Fig4; all data in Supp Table 1). To begin to determine how each cluster might be affecting RQDM, we focused on testing whether they might affect the electron transport chain directly since we previously showed that Complex I inhibitors or Complex II inhibitors are potent blockers of RQDM^11^. Using established individual *in vitro* assays for each of the four core complexes of the ETC, we confirmed that purified *C. elegans* mitochondria are specifically affected by known inhibitors (e.g. rotenone only affects Complex I activity, KCN only affects Complex IV) as shown in Supp Fig 1. We then tested members of each of the 7 clusters for ability to inhibit each of the individual complexes (results in Table 1). Several of the NP clusters appear to affect more than one Complex — for example anacardic acid is a potent inhibitor of Complexes III and IV, and Cluster 6 affects I, III, and IV. Other clusters (e.g. Cluster 2 and 3) have no consistent effect on any specific complex suggesting they likely have a different mode of action. However, Cluster 4 showed highly specific inhibition of Complex I. Complex I is the sole entry point for electrons into the ETC during RQDM and we previously showed that rotenone, a Complex I inhibitor, can block RQDM. We thus focused on characterising the compounds in Cluster 4.

**Figure 4:**
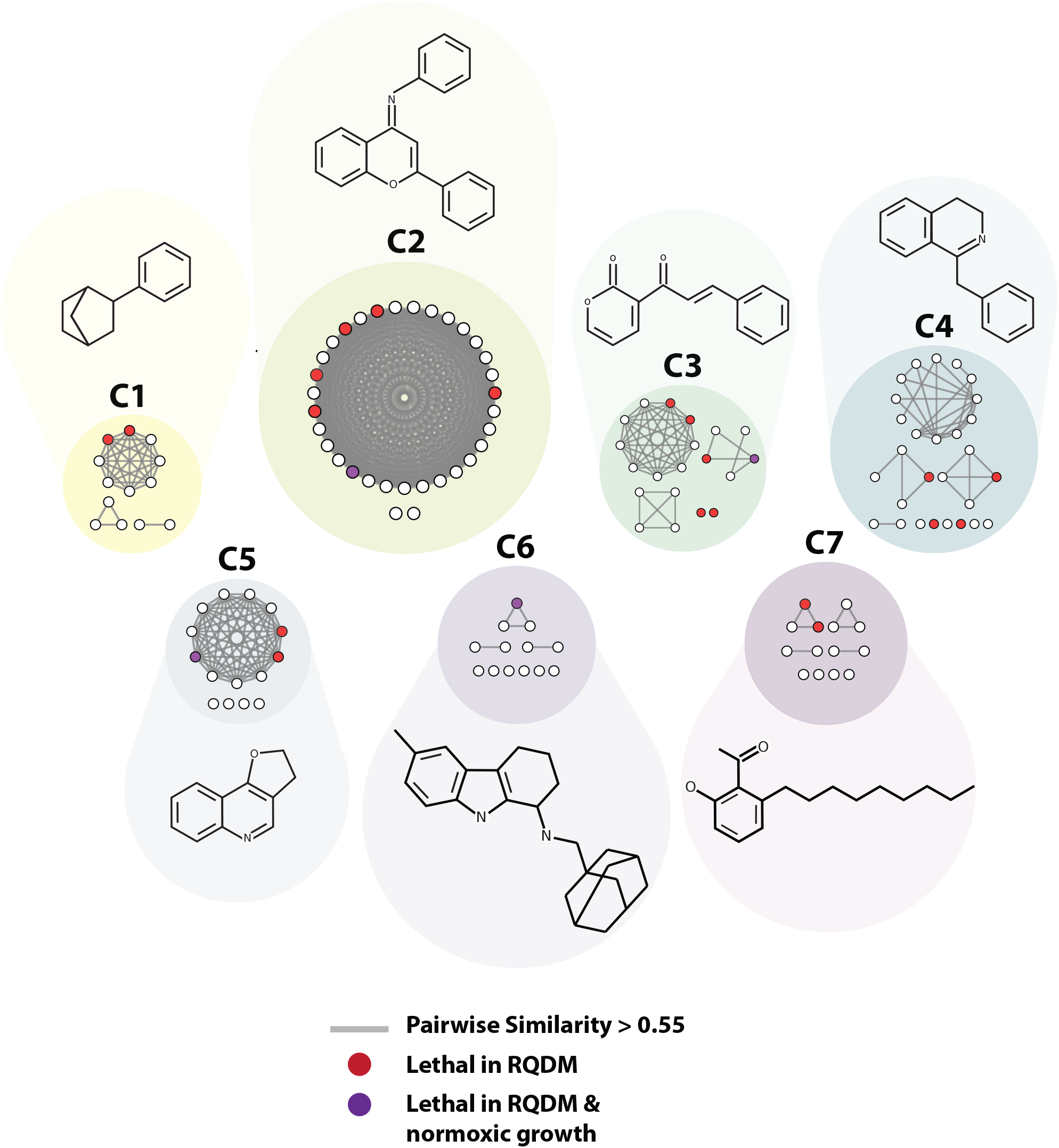
Structural Profiling of hits and associated derivatives/analogs from RIKEN NP library. Network based on the structural similarity of the 137 molecules from the seven structural classes identified as hits in the RIKEN NP Library. Nodes represent molecules, and edges connect molecules with scaffolds that have a pairwise Tanimoto/FP2 score > 0.55. The cluster to which each molecule belongs is indicated by the background circle, while the molecules effect in either condition of the screen is indicated by the node fill colour. The seven clusters are based off the chemical similarity performed by RIKEN NPDepo. Structures that share the similar scaffold are connected, while more distantly related molecule are related by a common substructure. Above each cluster is the murcko scaffold (for ring systems > 1) of the initial hit identified in the chemical screen.

**Table 1.** Summary of the effect of Riken NPDepo hits on the activity of mitochondrial ETC complexes (I-IV). Compounds that reduced activity of a complex below 50% at 100 μM are highlighted in red.

| Cluster | Compound | % Activity |  |  |  |
| --- | --- | --- | --- | --- | --- |
|  |  | I | II | III | IV |
| Control | Rotenone | 0.0 ± 0.0 | 97.7 ± 1.4 | 97.6 ± 1.8 | 97.9 ± 1.3 |
| Control | Wact-11 | 84.8 ± 2.9 | 24.3 ± 1.2 | 90.1 ± 3.5 | 89.0 ± 1.3 |
| Control | Antimycin A | 100 ± 0.0 | 75.9 ± 8.4 | 0.0 ± 0.0 | 94.0 ± 2.1 |
| Control | KCN | 100 ± 0.0 | 93.7 ± 6.3 | 98.1 ± 1.5 | 0.4 ± 0.4 |
| 1 | NPD8902 | 27.8 ± 1.4 | 100 ± 7.3 | 74.5 ± 5.5 | 0.8 ± 0.8 |
| 2 | NPD8034 | 99.3 ± 5.2 | 89.0 ± 9.5 | 95.1 ± 3.0 | 96.0 ± 5.6 |
| 2 | NPD1450 | 84.5 ± 4.1 | 100 ± 8.7 | 100 ± 1.9 | 100 ± 6.6 |
| 2 | NPD6380 | 86.4 ± 4.5 | 100 ± 4.9 | 100 ± 2.7 | 100 ± 8.8 |
| 3 | NPD8298 | 79.9 ± 2.5 | 100 ± 10.1 | 100 ± 4.9 | 100 ± 4.3 |
| 3 | NPD6240 | 100 ± 3.6 | 87.7 ± 7.6 | 58.2 ± 4.6 | 26.8 ± 1.5 |
| 3 | FSL0005 | 91.9 ± 1.5 | 80.9 ± 4.0 | 100 ± 2.5 | 92.3 ± 6.0 |
| 3 | NPD8366 | 100 ± 3.2 | 86.4 ± 6.7 | 76.0 ± 2.2 | 97.7 ± 5.5 |
| 4 | NPD10504 | 5.6 ± 3.0 | 100 ± 13.7 | 80.0 ± 3.0 | 74.5 ± 2.7 |
| 4 | NPD3577 | 36.8 ± 4.3 | 100 ± 9.9 | 66.0 ± 7.3 | 81.1 ± 4.7 |
| 4 | NPD6303 | 12.7 ± 1.3 | 100 ± 7.9 | 78.1 ± 2.8 | 83.1 ± 3.5 |
| 4 | NPD8790 | 5.3 ± 2.9 | 99.7 ± 8.6 | 64.0 ± 1.8 | 79.8 ± 2.7 |
| 5 | NPD390 | 18.5 ± 1.8 | 82.9 ± 3.7 | 85.2 ± 3.7 | 92.1 ± 4.2 |
| 6 | NPL50604-01 | 0.5 ± 0.5 | 82.6 ± 6.5 | 34.1 ± 0.6 | 21.5 ± 2.7 |
| 7 | Anacardic Acid | 93.0 ± 3.3 | 87.6 ± 2.7 | 0.0 ± 0.0 | 15.8 ± 2.3 |

We had identified 4 potent inhibitors of RQDM in cluster 4, shown in Fig 5a. None of these had been previously characterised but a search for structurally-related commercial compounds identified papaverine as a close analogue. Papaverine is largely reported to be an inhibitor of phosphodiesterase^41^ but there are 2 reports that it can also act as a Complex I inhibitor^42,43^ and we confirm that it acts as a Complex I inhibitor in our hands on purified *C. elegans* mitochondria (Fig 5b). We next wanted to test whether any of the cluster 4 hits, or papaverine itself, showed any species selectivity — Complex I is critical for oxidative phosphorylation in the vertebrate hosts and thus we ideally want a compound that can selectively target the helminth Complex I. We found that neither papaverine nor three highly similar NP hits showed any significant species selectivity (Fig 5c and data not shown). However, NPD8790 showed good species selectivity, having >10-fold difference in IC50 between *C. elegans* Complex I and either bovine or murine Complex I (Fig 5d). NPD8790 is structurally distinct from the other compounds in cluster 4 — instead of a quinoline or isoquinoline core, the core of NPD8790 is a benzimidazole. This is intriguing since benzimidazoles (BZs from here on) are extremely well characterised as anthelmintics and include the front line anthelmintics mebendazole, albendazole and related drugs^19,20^. The mode of action of known anthelmintic BZs is well established: they target beta-tubulins^14,16,17,19^ and affect their ability to polymerise. We therefore wanted to examine whether known BZs also affect Complex I and whether NPD8790, the BZ that we identified as a Complex I inhibitor, also targets BEN-1.

**Figure 5:**
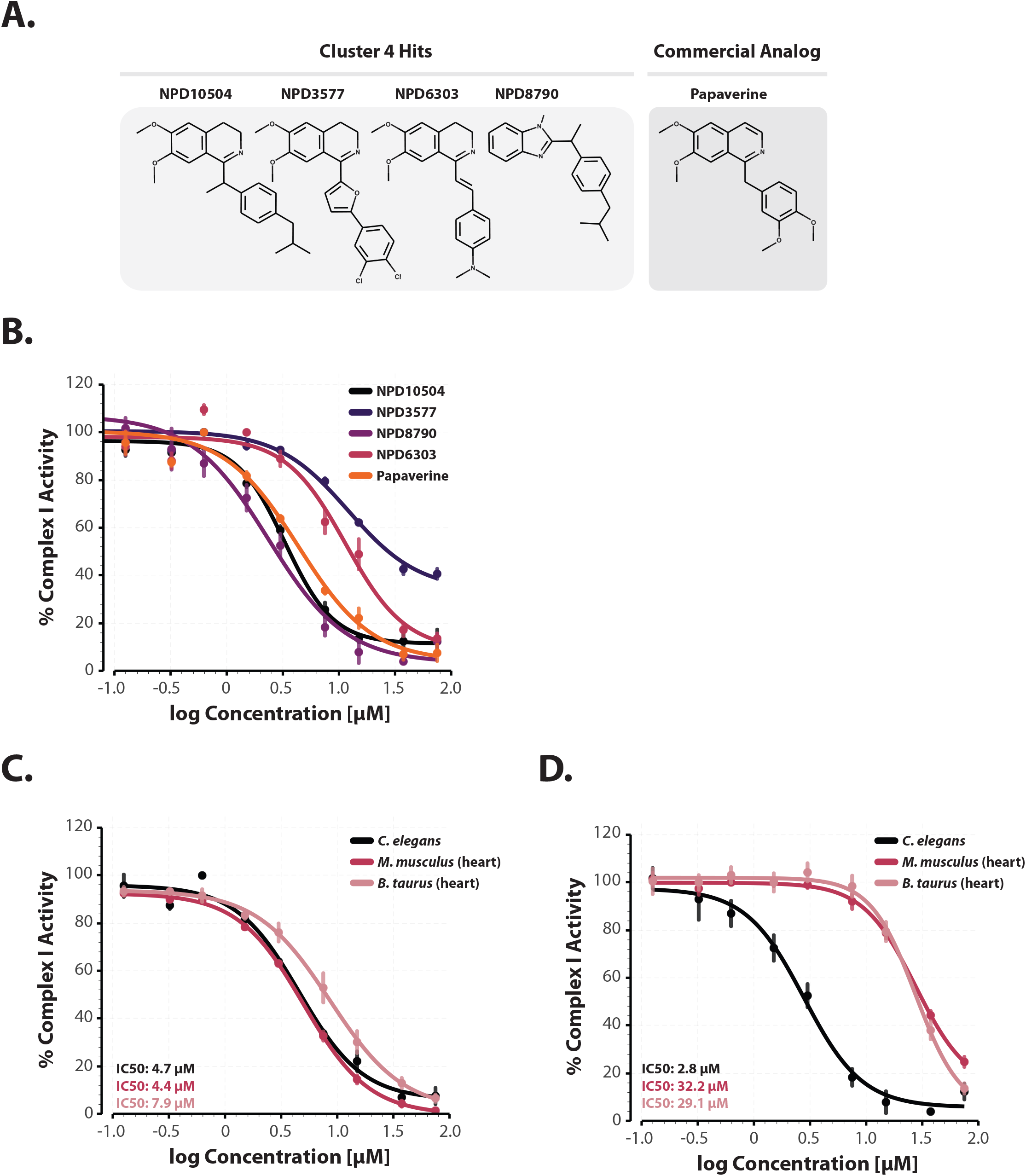
Cluster 4 compounds inhibit Complex I of the ETC. A. Structures of Cluster 4 hits and papaverine. **B. Papaverine and structurally related cluster 4 compounds inhibit Complex I**. Complex I activity was assessed spectrophotometrically at 600nm following reduction of a colorimetric dye (DCIP) by purified *C. elegans* (N2) mitochondria. Data are mean of 4 independent replicates; error bars show the standard error of the mean. All four cluster 4 compounds inhibit complex I like papaverine, an FDA approved spasmodic with documented effects on metabolism and complex I inhibition. **C. Papaverine shows no species selectivity.** Purified mitochondria from *C.elegans*, mouse heart or bovine heart were treated with different doses of papaverine and Complex I activity measured. **D. NPD8790 shows significant species selectivity.** Purified mitochondria from *C.elegans*, mouse heart or bovine heart were treated with different doses of NPD8790 and Complex I activity measured.

### Commercial benzimidazole anthelmintics do not target Complex I

In *C. elegans*, the principal target of commercial anthelmintic BZ drugs is the beta-tubulin BEN-1^14^. However, we found that NPD8790 is a potent inhibitor of Complex I and thus we wanted to test whether commercial anthelmintic BZ drugs also affect Complex I activity. We find that none of the commercial BZs have any detectable effect on Complex I activity in purified mitochondria suggesting NPD8790 has a new activity for BZ compounds (Fig 6). In *C. elegans*, mutation or deletion of BEN-1^44,45^ results in resistance to commercial anthelmintic BZs. If NPD8790 also targets BEN-1, it should be similarly affected by BEN-1 mutation or loss. We confirmed that in our hands, multiple commercially available BZ anthelmintics potently kill *C. elegans* in normoxia (Fig 7A and Supp Fig 2) and that either deletion of *ben-1* or any of the *ben-1* mutations known to cause BZ resistance in parasites^16,17^ greatly reduce sensitivity as expected (Fig 7A). However, NPD8790 has very little effect on *C. elegans* normoxic growth until at very high concentrations and the small effect on growth is unaffected by *ben-1* mutation suggesting it may be acting in a different manner (Fig 7C). We also find that commercial BZ anthelmintics have little effect in our RQDM assay (Fig 7B and Supp Fig 2) and that the inhibition of RQDM by NPD8790 is not affected by mutations in *ben-1* (Fig 7D). Finally, we purified mitochondria from several *C. elegans* strains carrying *ben-1* mutations and tested whether NPD8790 could still affect Complex I activity. We find that mutations in *ben-1* have no effect on the ability of NPD8790 to inhibit Complex I (Fig 7E). We thus conclude that NPD8790 is a BZ compound that has a distinct mode of action to all available BZ anthelmintics — while they target BEN-1 and not Complex I, NPD8790 targets Complex I and not BEN-1.

**Figure 6:**
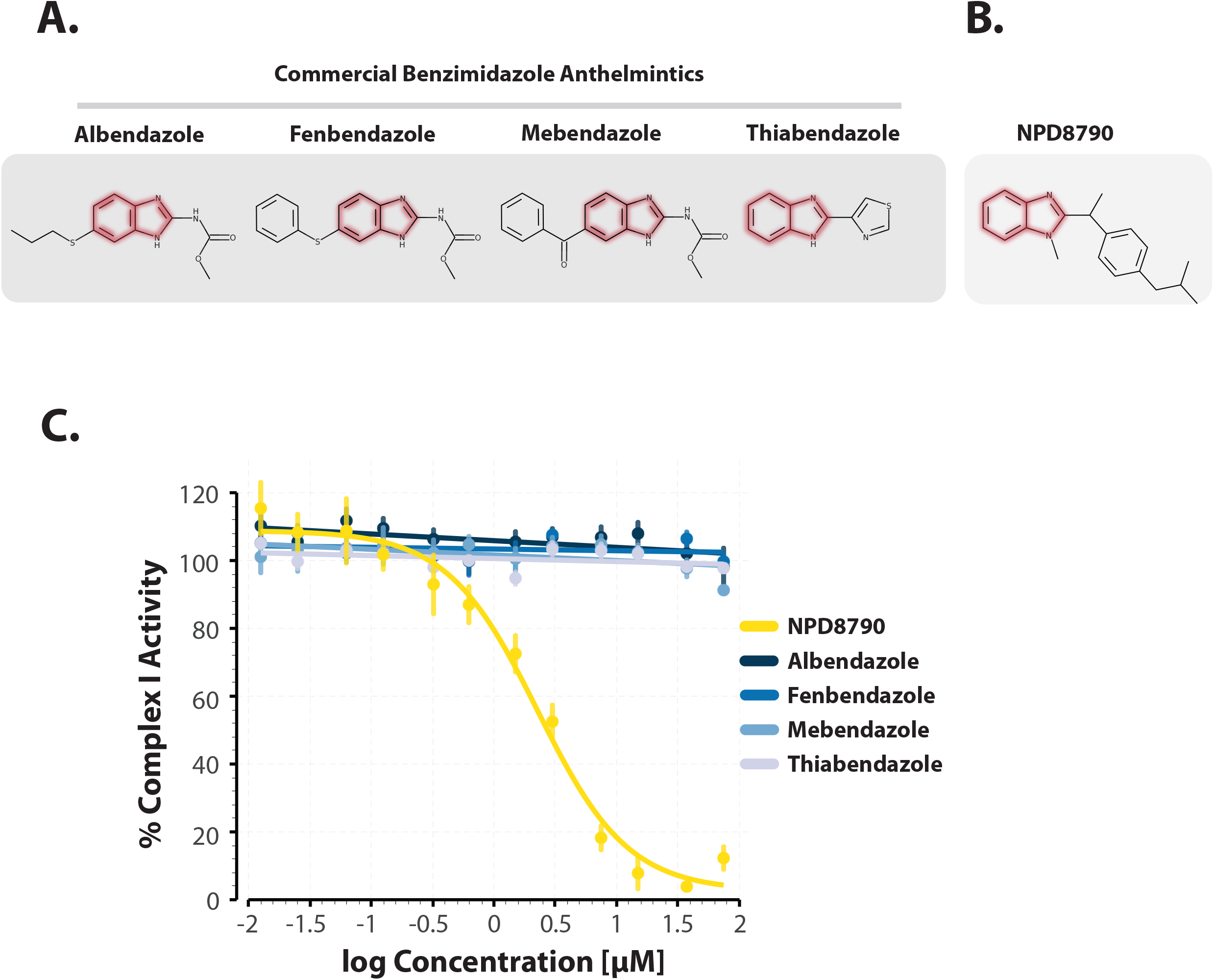
NPD8790 affects Complex I activity but commercial benzimidazoles do not. **A-B.** Structures of 4 benzimidazole anthelmintics and NPD8790. **C.** Complex I inhibitory dose-response curves for NPD8790 and several commercial benzimidazole anthelmintics (Albendazole, Fenbendazole, Mebendazole, Thiabendazole) against complex I from wild-type (N2) *C. elegans.*

**Figure 7.**
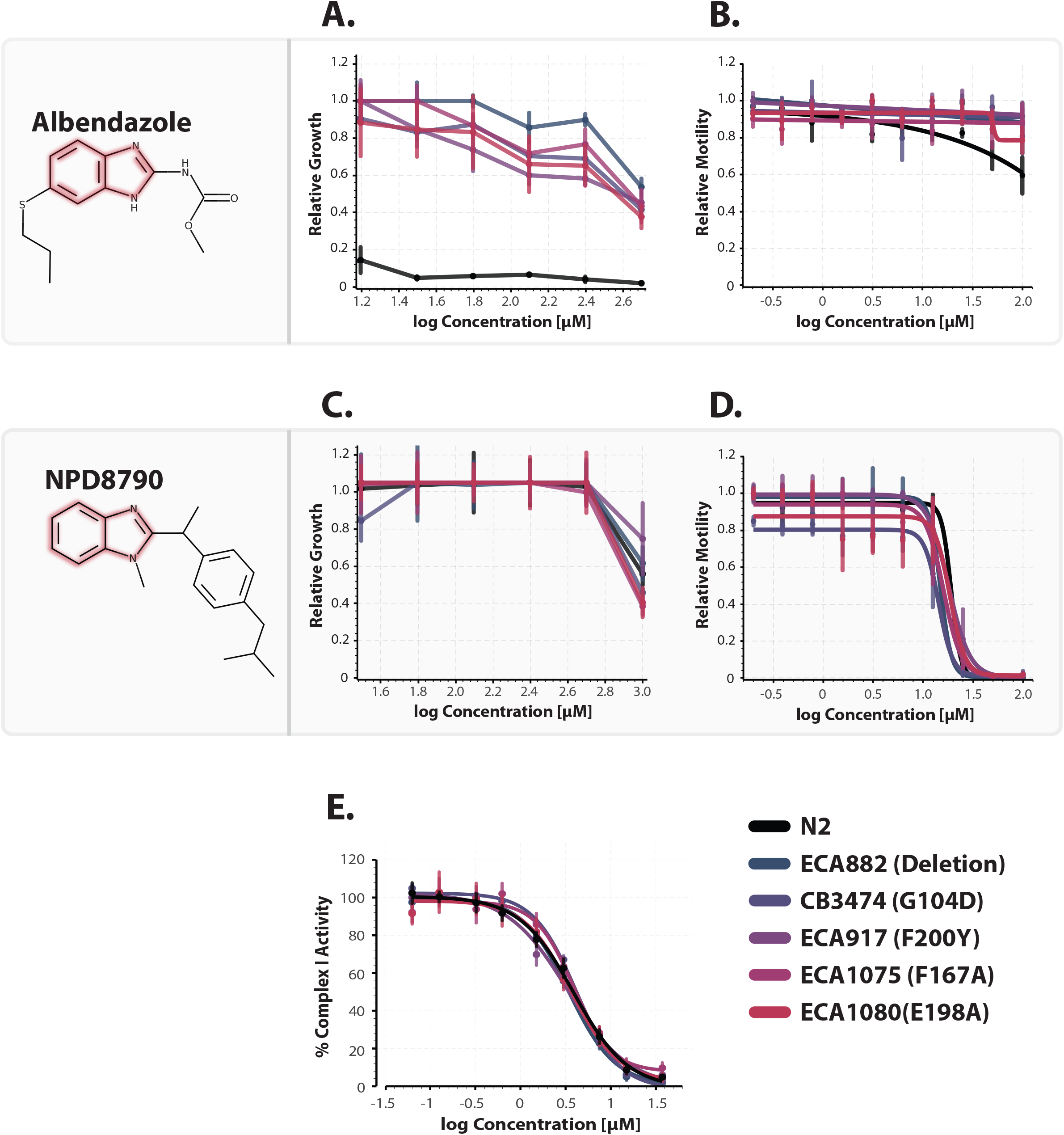
Loss of function mutations in the *C. elegans ben-1* gene confer a high degree of resistance to commercial benzimidazole anthelmintics, but not to NPD8790. **A. Inhibition of worm growth by albendazole is affected by *ben-1* mutations**. *C. elegans* strains were exposed to different doses of albendazole and the growth was measured after 6-days in the presence of drug. Data are mean of 4 independent replicates; error bars show the standard error of the mean. Wild-type (N2; black) worms were strongly affected but strains homozygous for different mutations in the *ben-1* gene show strong resistance. **B. Albendazole has no effect on RQDM.** *C. elegans* strains were exposed to different doses of albendazole in the presence of 200 μM KCN for 15hrs. Drugs were diluted and movement was measured after 3hrs. Wild-type (N2; black) and strains carrying mutations in *ben-1* all showed good recovery of movement. **C. NPD8790 has little effect of *C.elegans* growth in normoxia.** *C.elegans* strains were exposed to different doses of NPD8790 and the growth was measured after 6-days in the presence of drug. Data are mean of 4 independent replicates; error bars show the standard error of the mean. Wild-type (N2; black) worms and strains homozygous for different mutations in the *ben-1* gene show no growth defects relative to controls except at extremely high NPD8790 concentrations. **D. NPD8790 blocks RQDM.** *C.elegans* strains were exposed to different doses of NPD8790 in the presence of 200 μM KCN for 15hrs. Drugs were then diluted, and movement was measured after 3hrs. Wild-type (N2; black) and strains carrying mutations in *ben-1* all showed similar inhibition of RQDM. Data are mean of at least 4 independent replicates; error bars show the standard error of the mean. **E. *ben-1* mutations have no effect on ability of NPD8790 to inhibit Complex I.** Mitochondria were purified from wild-type worms or from strains carrying a variety of *ben-1* mutations and Complex I activity assayed at a range of doses of NPD8790.

### Structure-function analysis identifies requirements for Complex I inhibition by NPD8790 BZ compounds

The sole BZ compound that we identified as a Complex I inhibitor in our NPD screens was NPD8790. To explore how it targets Complex I and to test whether we could identify related compounds with either higher potency or higher species selectivity, we screened 1286 structurally related BZ compounds. We first assayed each compound in our RQDM assay at 50*μ*M to narrow down BZ compounds that potentially shared similar bioactivity with NPD8790. A total of 88 BZ compounds (6.8%) reduced motility by at least 75% in our RQDM assay, 64 of these compounds that were available were then reordered to perform more comprehensive dose responses. Each of the 64 compounds were assayed for their effect on Complex I activity in both *C. elegans* and bovine mitochondria, as well as their ability to affect *C. elegans* growth. Finally, we also assayed toxicity in HEK293 cells. These data are all shown in Supp table 2.

We identified 3 distinct structural classes of BZ compounds that are Complex I inhibitors that block RQDM in our assays and that show potential as anthelmintics — we will refer to these as classes A-C (Fig 8). All of them have a BZ core which is linked to an aromatic ring. The key differences between the subclasses is the position and the length of the linkage. Class A compounds include NPD8790 and in all of these the aromatic ring is linked via position 2 of the BZ group via a short linkage (1-atom linkage). These compounds are good Complex I inhibitors with IC50s well below 10 μM. More importantly, Class A includes the compounds with the strongest species selectivity e.g. NPD8790 has a >10-fold differences in IC50 between *C. elegans* and bovine Complex I. They have very little detectable effect on growth in normoxic conditions and NPD8790 also has no observable toxicity in mammalian cells even at high concentrations (>50 μM). These are thus good candidates as anthelmintics.

**Figure 8.**
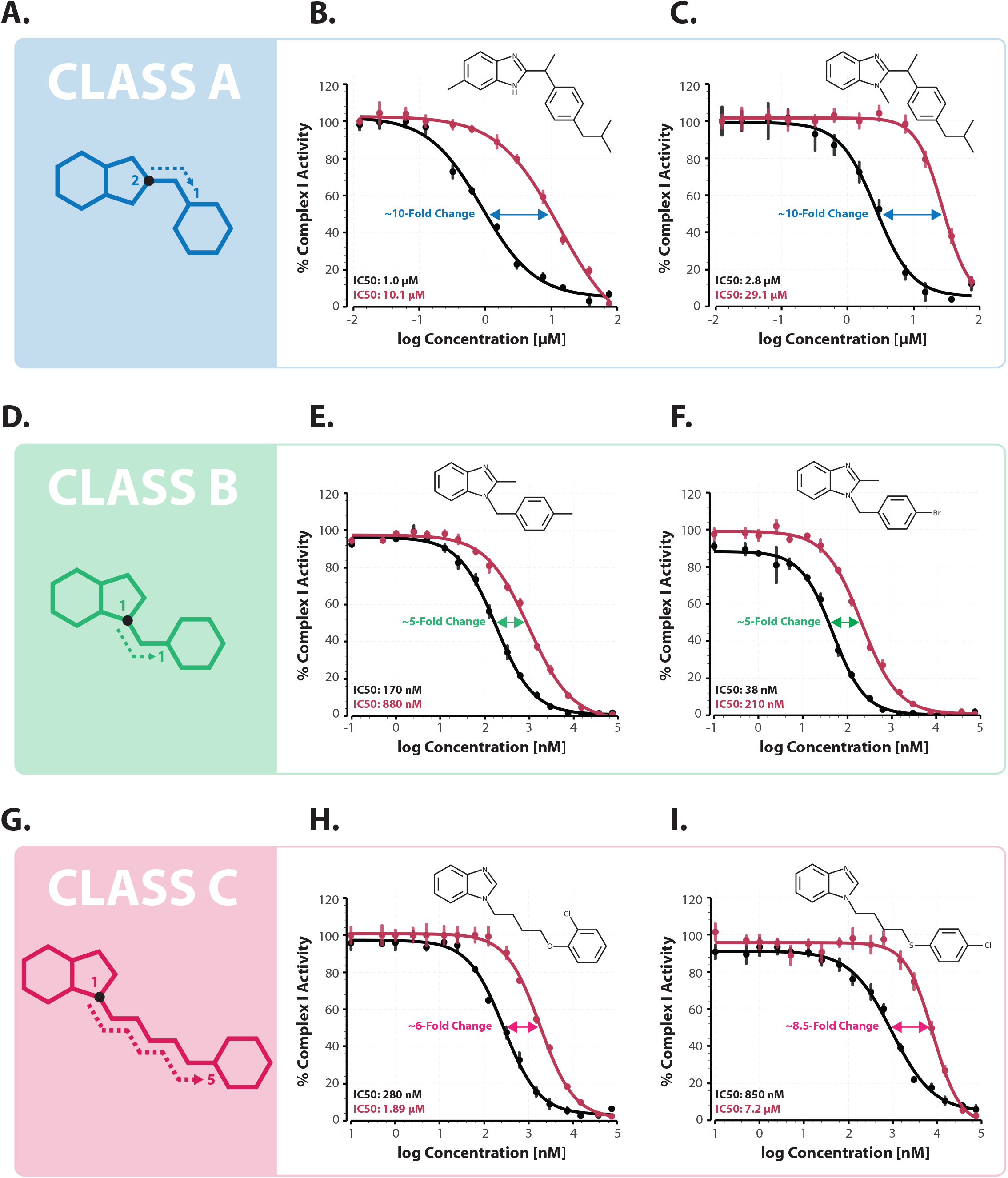
Three different classes of benzimidazoles act as Complex I inhibitors. Panels **A, D** and **G** show core structures of Class A, B and C benzimidazoles identified as Complex I inhibitors in our screens. In the remaining panels we show inhibition of Complex I activity either in purified *C. elegans* (black curves) or bovine mitochondria (red curves) for representative molecules from each Class. Each curve is the mean of 2 independent replicates; errors are the S.E.M.

The other two classes both contain multiple compounds with substantially more potent Complex I inhibition than Class A e.g. Chembridge ID# 5567167 has a 75-fold lower IC50 for *C. elegans* Complex I than NPD8790. These compounds all have the aromatic rings linked through the 1st position of the BZ core with linkages of varying lengths — the differences between the two classes is in the length of the linkage between the aromatic group and the BZ core. Class B comprises compounds with a 1-atom linkage — these are very potent inhibitors with reasonable species selectivity. Compounds with 2-atom linkages are substantially less potent, but potency appears to improve with longer linkages. 5-linkage compounds show excellent potency with good species selectivity, and these make up Class C of our potential anthelmintics. We note that Class B and C compounds are good anthelmintic candidates but that both sets show effects on growth in normoxia unlike Class A compounds, they also show a slightly reduced species selectivity. Consistent with their effect as Complex I inhibitors, we find that treatment of wild-type worms with the Class B and C BZ compounds causes a rapid decrease in ATP levels (Supp Fig 3) and a rapid drop in mitochondrial potential. This effect on ATP levels in normoxia is also seen for Class A compounds but at much higher concentrations. This underlies that one of the major differences between Class A and Class B/C is the effect in normoxia. This suggests that Class A compounds may selectively effect worms when they rely on RQDM, making them good candidates for anthelmintics.

## Discussion

Soil-transmitted helminths (STHs) are major human pathogens. While there are excellent frontline anthelmintics including macrocyclic lactones and benzimidazoles, the number of classes of anthelmintics is very limited and resistance is increasing and is already very high for some livestock parasites. There is thus an urgent need for new classes of anthelmintic drugs. In this study we focused on using *C. elegans* to screen for drugs that specifically inhibit the unusual anaerobic metabolism that STHs like *Ascaris* and hookworm use to survive the low oxygen conditions in the host gut. This metabolism requires rhodoquinone (RQ), an electron carrier that is absent from the hosts. We screened a library of natural products and their derivatives (NPDs) that contained both previously characterised and uncharacterised compounds and identified multiple distinct structural families of compounds that kill *C. elegans* specifically when they are surviving using RQDM — we term these ‘RQDM hits’. These are candidate anthelmintics since they may similarly kill STHs in vivo when they rely on RQDM for survival in the gut.

We assessed whether the RQDM hits target specific complexes of the ETC. During RQDM, electrons enter the ETC solely through Complex I and exit through Complex II and consistent with this we had previously shown that known Complex I inhibitors and Complex II inhibitors are potent inhibitors of RQDM. We identify both specific Complex I and Complex II inhibitors in our screens — a novel set of benzimidazole compounds as Complex I inhibitors and Siccanin as a Complex II inhibitor. The way these two sets of compounds affect RQDM is consistent with our previous findings as these are expected to block electron entry or exit from the ETC. Other RQDM hits are more enigmatic however. For example, anacardic acid is a potent inhibitor of Complex III and IV in purified mitochondria and it is unclear mechanistically how it might be acting or why it has such a strong effect on RQDM but appears to have no detectable effect on metabolism in normoxia. Other RQDM hits appear to have no effect on the ETC and must be affecting RQDM in some other unknown manner, perhaps through targeting other required components of RQDM such as AMPK^46^ or on RQ synthesis itself. Thus while the mode of action of some of the RQDM hits fits with what is known of electron flow in the ETC in RQDM, others will require future studies to identify the targets and mode of action.

The main focus of our analysis was on a novel family of benzimidazole (BZ) compounds that we identified as potent and specific Complex I inhibitors. We showed that the effect of these new BZ compounds is not altered by mutations in *ben-1*, the beta tubulin target of well-characterised commercial benzimidazoles, and that commercial benzimidazoles do not affect Complex I activity confirming that the inhibition of Complex I by this new family of BZ compounds is a novel activity.

The critical difference between the new BZs we identified as RQDM hits and BEN-1-targeting commercial BZ drugs is the linkage of a benzene ring to the benzimidazole ring in the new Complex I inhibiting BZ compounds. The linkage length and precise position of the linkage on the benzimidazole ring affects potency and specificity in complex ways. We find that the most potent Complex I inhibitors have the benzene ring attached to the 1-position of the benzimidazole ring with a 1-atom linkage (Class B) — the most potent such compounds have ~100nM IC50s for Complex I. However, they affect both bovine and *C. elegans* Complex I with similar IC50s, suggesting that they are not ideal as anthelmintics. Compounds with the same 1-atom linkage between the benzimidazole and benzene group but linked to the 2-position of the benzimidazole ring show slightly reduced inhibition of Complex I but increased species selectivity and essentially no detectable effect on growth in normoxia when the worms are using Complex I coupled to ubiquinone. The difference in the effect of these compounds on ubiquinone-coupled electron transport in normoxia and on RQ-coupled electron transport in KCN conditions is marked — for example NPD8790 has no effect on growth in normoxia at 1 mM but has an LD50 in our RQDM assay of ~14 μM. We speculate that this may reflect a difference in ability to block docking of UQ and RQ to Complex I and that this may also underlie the species selectivity we observe. This set of compounds where the aromatic group is linked via a 1-atom linkage on the 2 position of the benzimidazole ring (Class A) appear like much better candidates as anthelmintics given their higher species selectivity, and the absence of any detectable effect on normoxic growth. These are our leading candidates as anthelmintics.

We also note that lengthening the linkage between the BZ core and the benzene ring has complex effects. Increasing from a 1-atom to a 2-atom linkage length greatly reduces the ability of the compounds to inhibit Complex I. For example, while compounds with the benzene group linked to the 1st position of the benzimidazole ring with a 1 atom linker (Class B) have IC50s in the sub micromolar range, this increases to >50μM when the linkage is a 2-atom linkage. Intriguingly, this dramatic drop in potency is highest for 2-atom linkages — 3- and 4-atom linkages to the same benzimidazole position have low μM IC50s for Complex I and finally compounds with 5-atom linkages (Class C) typically show excellent potency with IC50s in the sub micromolar range like those with a 1-atom linkage (Class B). This high potency with either a 1 or 5-atom linkage and reduced potency with 2-4 atom linkages is intriguing. We suggest that the 1-atom and 5-atom linkage compounds are interacting with the active site in distinct ways and speculate that this may reflect the complexity of the way that Complex I interacts with quinones through its multiple Fe-S centres. Finally, we note that there is a single previous report of an insecticide with a benzimidazole core with activity as a Complex I inhibitor^47^. However, the structure is distinct to the hits we have identified here — instead of an aromatic group linked to the BZ core, it has a long terpenoid chain. Additionally, no species selectivity has yet to be demonstrated for this compound.

In summary, we screened a natural product library in *C. elegans* to identify compounds that killed *C. elegans* only under conditions when they require RQDM to survive. We believe that this is a reasonable model for helminths which rely on RQDM to survive in the highly anaerobic environment of a host gut. We identified compounds that specifically inhibit either Complex I or Complex II in a species-specific manner. Complex I and II are the entry and exit points for electrons in the RQ-coupled ETC and thus the mode of action is consistent with what is known of the ETC in RQDM. The Complex I inhibitors are a family of compounds with a benzimidazole core and thus is a novel activity for benzimidazoles — well characterised benzimidazole anthelmintics have no activity against Complex I and target microtubules. We thus believe that we have a new class of compounds as potential anthelmintics that act by specifically targeting the unusual anaerobic metabolism of this major class of animal and human pathogens.

## Acknowledgements

This research was funded by grants 501584 and 5003009 to Andrew Fraser from the Canadian Institute of Health Research. We thank Prof Peter Roy and Prof Erik Andersen for sharing worm strains; Prof Derek van der Kooy, Brenda Takabe and Daniel Merritt for mouse organs and use of their Dounce homogenizer; and the Peel Sausage company for the beef heart. We also thank Jason Moffat’s group for letting us use their plate reader. Finally, we thank Andrew Burns and Sam Del Borrello for guidance and helpful discussion throughout.

## Materials and Methods

### Chemical Sources

The Authentic and Pilot libraries of natural products as well as related derivatives used in our preliminary screens were provided by the RIKEN Natural Product Depository (NPDepo). Hit compounds identified were purchased from Vitas-M for retesting. Benzimidazole analogs of NPD8790 were purchased from ChemBridge Corporation, Vitas-M and Enamine. Wact-11 (ID # 6222549) was purchased from ChemBridge Corporation. Rotenone, antimycin A, potassium cyanide, albendazole, mebendazole, thiabendazole, fenbendazole were purchased from MilliporeSigma. Siccanin was purchased from Cayman chemical.

### *C. elegans* strains and maintenance

All animals were cultured using standard methods at 20°C on NGM (nematode growth medium) agar plates seeded with E. coli OP50 as previously described (Stiernagle, 2006)^48^. In addition to the traditional laboratory strain N2, this work includes the following strains: CB1003 (*kynu-1(e1003)*), RP2639 (*sdhc-1(tr359)*), RP2702 (*sdhc-1(tr410)*), RP2748 (*sdhc-1(tr4230)*), RP2749 (*sdhc-1(tr424)*), RP2776 (*sdhb-1(tr438)*), CB3474 (*ben-1(e1880)*), ECA882 (*ben-1(ean64)*), ECA917 (*ben-1(ean98)*), ECA1075 (*ben-1(ean143)), ECA1080 (ben-1(ean148)*), and PE255 (feIs5 [*sur-5*p::luciferase::GFP + *rol-6(su1006)]* X). The five *sdhc-1* and *sdhb-1* strains were generously provided by Dr. Peter Roy, five *ben-1* strains (ECA prefix) were generously provided by Dr. Erik Andersen, and all other strains were provided by the *Caenorhabditis* Genetics Centre (CGC, University of Minnesota).

### Image-based chemical screens in *C. elegans*

The rhodoquinone-dependent metabolism (RQDM) assays were performed as described previously (Del Borrello *et al.*, 2019) using an image-based system for measuring worm motility that was developed in lab (Spensley *et al.*, 2018)^49^. Briefly, 25 *μ*L of M9 buffer was distributed to each well of a flat-bottomed 96-well culture plate, and chemicals were added using a pinning tool with a 500 nL slot volume (V&P Scientific). First larval-stage (L1) worms were isolated from mixed-stage plates using 96-well 11 *μ*M nylon mesh filter plates (Millipore Multiscreen) and diluted in M9 to a concentration of ~6 worms/μL. 20 *μ*L of worm suspension containing approximately 120 animals was then added to each well. Stock solutions of potassium cyanide (KCN) were prepared fresh in phosphate buffered saline (PBS) before each experiment and diluted to a 10X working concentration (2mM) in M9 buffer. 5 *μ*L of 10X KCN solution was added to each well to reach a final concentration of 200 *μ*M KCN. Immediately following the addition of KCN, plates were sealed with aluminum sealing tape to prevent evaporation and were incubated for 15 hours at 20°C with shaking at 165 rpm. Following 15 hours of incubation, KCN in wells was diluted 6-fold with M9 buffer and plates were placed on a Nikon Ti Eclipse inverted microscope to be monitored every 10 minutes for 3 hours. For each timepoint, two images of every well were captured at an interval of 500 ms and processed with custom Python scripts to isolate worm-associated pixels from background. An aggregate score of raw worm motility was determined for each well by comparing the change in worm-associated pixels between the two images (i.e., worm movement between two frames) to the sum of all worm-associated pixels (i.e., number of worms in frame). The raw motility score for each well was divided by the raw motility scores for the corresponding dimethyl sulfoxide (DMSO) controls, resulting in a “Relative motility” value for each chemical. For chemical screens, the relative motility of each well after 3 hours of KCN dilution was used as an endpoint.

The *C. elegans* liquid growth and viability assays were adapted from a previously established chemical screening protocol (Burns *et al.*, 2015)^39^. In short, a saturated culture of HB101 *E. coli* was concentrated 2-fold in liquid NGM and 40 *μ*L of this suspension was dispensed to each well of a flat-bottomed 96-well culture plate. Chemicals from each library were added using a pinning tool with a 500 nL slot volume (V&P Scientific). L1 worms were synchronized from an embryo preparation (Burns *et al.*, 2006)^40^ performed the day prior and diluted to a concentration of ~2 worms/μL in M9 buffer. 10 μL of worm suspension containing approximately 20 animals was then added to each well. Culture plates were sealed with parafilm and placed in a plastic container lined with wet paper towel to prevent evaporation. Containers were then transferred to incubators for 6 days at 20°C with shaking at 165 rpm. Following incubation, culture plates were removed, and wells were diluted 7-fold with M9 to decrease the turbidity of the HB101-NGM suspension. 1-2 minutes post-dilution, culture plates were placed on a Nikon Ti Eclipse inverted microscope and single images were taken of each well. Images were processed using custom Python scripts to isolate worm associated pixels from background. An overview of the image processing steps has been previous described (Spensley *et al.*, 2018)^49^. Raw scores of overall worm growth and viability were determined as the sum of worm-associated pixels for each well. The raw growth/viability score for each well was divided by the raw grow/viability scores for the corresponding DMSO controls, resulted in a “Relative growth” value for each chemical.

NP libraries were stored in 96-well plates, each containing 80 compounds and 16 DMSO controls (columns 1 and 12). For our preliminary screens of NP libraries, six 96-well plates containing 480 compounds and 96 DMSO controls were screened in triplicate at a concentration of 50 μM for both *C. elegans* assays; data is displayed as the average of the three biological replicates. The final concentration of DMSO in wells was kept to 1% v/v to prevent any confounding effects of drug solvent on worm motility or viability. To account for possible variation among plates, the process of hit identification was performed on a plate-to-plate basis. As drug hits (i.e., outliers) can influence the mean of the distribution of scores, relative motility or relative growth values were converted to robust z-scores (r.z-score) using the median and the median absolute deviation (MAD) of each plate. Hits were identified if a compound’s r.z score fell below −3 (i.e., 3x MAD below the median) in either of the respective assays.

Each of these assays was used for additional follow-up dose-response or time course experiments in this work. These experiments were carried out with at least three or more biological replicates. For dose-response experiments, dose response curves were fitted using non-linear regression in the Python library Scipy. LC50 and EC50 values were obtained from the fitted curves.

### HEK293 cell culture and dose-response experiments

Human embryonic kidney (HEK293; Invitrogen’s Flp-In-293 cell line cat #R75007) cells were grown and maintained in Dulbecco’s Modified Eagle’s Medium (DMEM) containing 10% fetal bovine serum (FBS) and 1% penicillin-streptomycin (PS). To prepare cells for viability testing, 5000 HEK293 cells/100 *μ*L were seeded into each well of a 96-well flat-bottom tissue culture plate and grown overnight at 37°C with 5% CO_2_. 0.5 *μ*L of compound from prepared dose-response plates were then added to each well (0.5% v/v DMSO) and cells were allowed to grow for an additional 2 days. After this period of growth, cells were incubated for 4 hours with 10 *μ*L of CellTiter-Blue viability reagent. Raw scores of cell viability were then determined using a CLARIOstar Plate Reader (560/590nm) and the fluorometric quantification of the reduction of resazurin to resorufin. “Relative viability” was determined by dividing the background-corrected scores of chemical-treated wells from the corresponding DMSO controls. Data represents the average of three biological replicates.

### Cheminformatics

Clusters of natural products and derivatives related to each of the initial hits identified in our chemical screen were curated by the RIKEN natural product depository. Chemical structures of these compounds were analyzed in DataWarrior (Sander *et al.*, 2015)^50^. Chemical similarity among compounds of the same cluster was determined by identifying compounds with a shared core scaffold. Murcko scaffolds (for compounds with ring systems > 1) were first identified for each molecule using DataWarrior. FP2 fingerprints were then generated for each scaffold using Pybel (O’Boyle *et al.*, 2008)^51^ and pairwise tanimoto coefficients between fingerprints were used to assess chemical similarity. Scaffolds that shared a tanimoto coefficient of 0.55 or greater were grouped together. Network visualization of similar scaffolds in each cluster of natural products in our screen are shown using Cytoscape (Cline *et al.*, 2007)^52^.

### Mitochondria isolation from *C. elegans*

Mitochondria were isolated from adult *C. elegans* for each of the following strains: N2, RP2749, CB3474, ECA882, ECA917, ECA1075, and ECA1080. To obtain mitochondria in sufficient quantities, worms were grown in bulk through liquid culture. The isolation of mitochondria followed the protocol previously outlined in Burns *et al.*, 2015. The procedure was repeated for each of the respective strains.

### Mitochondria isolation from mammalian tissue

Six 8–10-week-old C57Bl/6 female mice (Charles River) were freshly dissected, and their livers and hearts collected into separate 1.5 mL microcentrifuge tubes. Organs were flash-frozen in liquid nitrogen and stored at −80°C prior to use. The heart of a freshly slaughtered adult cow was obtained from a local abattoir (Peel Sausage Inc), chopped into ~1-inch cubes and collected into 50 mL falcon tubes. Falcon tubes were flash frozen on location in a container of dry ice and later transferred to storage at −80°C to await mitochondria isolation. Isolation of mitochondria from all mammalian tissue was carried out following the protocol previously described in Burns *et al.*, 2015. However, due to size, the initial homogenization of cow heart tissue was performed with several 5 s pulses in small Nutribullet blender rather than using a Dounce homogenizer. All mouse protocols were reviewed and approved by the University of Toronto Animal Care Committee, in accordance with the Canadian Council on Animal Care.

### Electron Transport Chain (ETC) Assays

The enzymatic activities of individual ETC complexes were determined spectrophotometrically in isolated mitochondria using a Varioskan LUX Multimode plate reader (ThermoFisher Scientific). Mitochondrial pellets previously isolated were thawed on ice and resuspended in ice-cold isolation buffer (250 mM sucrose, 10 mM Tris (pH 7.5), 1 mM EDTA). The BCA assay (Walker, 1994) was used to quantify the protein concentration of mitochondria suspensions. Solutions were diluted to a concentration of 0.2 mg ml^-1^ for use in all assays. For use in the complex I assay, mitochondria pellets were freeze-thawed three times to increase accessibility to enzyme.

Rotenone-sensitive complex I (NADH: Decylubiquinone Oxidoreductase) activity was assessed using the 2,6-dichlorophenolindophenol (DCIP)-coupled method previously optimized in Long *et al.*, 2008^53^. Assays were setup in 96-well flat bottom culture plates, with each well containing 100 μM of chemical dissolved in 180 μL of complex I assay buffer (25 mM KPi buffer (pH 7.5), 3 mg mL^-1^ bovine serum albumin (BSA), 80 μM NADH, 60 μM decylubiquinone, 160 μM DCIP, 2 μM antimycin A and 2 mM KCN). DMSO (2.4% v/v) and rotenone (10 μM) controls were included in parallel to account for confounding effects of the solvent and determine the contribution of any rotenone-insensitive activity. The reaction was initiated by pipetting 20 μL mitochondria suspension into each well and briefly mixing. Absorbance was measured at 600 nm in 30 s intervals over the course of 15 minutes. Total complex I enzymatic activity was determined by plotting absorbance versus time for each well and calculating the slope of the line during the linear phase of the initial rate of reaction (minutes 1-4). Rotenone-sensitive activity was calculated by subtracting the complex I activity of the rotenone control wells. Percent complex I activity for each chemical was then calculated by dividing the rotenone-sensitive activity of chemical-treated wells by that of the DMSO control wells.

Complex II (Succinate Dehydrogenase) activity was assessed using the DCIP-coupled method previously described in Burns *et al.*, 2015^39^. Assays were similarly setup in 96-well culture plates, with each well containing 100 μM of chemical dissolved in 150 μL of complex II assay buffer (1X PBS, 0.35% BSA, 20 mM succinate, 240 μM KCN, 60 μM DCIP, 70 μM decylubiquinone, 25 μM antimycin A, 2 μM rotenone). DMSO, water and malonate (100 mM) controls were included in parallel. The reaction was initiated by pipetting 5 μL of mitochondria suspension into each well and briefly mixing. Absorbance was measured at 600 nm in 30 s intervals over the course of 15 minutes. Complex II enzymatic activity was determined by plotting absorbance versus time for each well and calculating the slope of the line during the linear phase of the initial rate of reaction (minutes 1-7). Percent complex II activity for each chemical was calculated by dividing the activity of chemical-treated wells by that of the DMSO controls.

Antimycin A-sensitive Complex III (Decylubiquinol-Cytochrome C Reductase) activity was determined by measuring the reduction of cytochrome C (CytC) in a protocol optimized in Luo *et al.*, 2008^54^. Decylubiquinol for complex III assays was prepared fresh as described in Janssen and Boyle, 2019^55^. In short, a few flakes of potassium borohydride were mixed into 10 mM decylubiquinone dissolved in ethanol, 0.1 M HCl was then added in 5 *μ*L increments until the solution turned colourless. The solution was spun down at 10,000 xg for 1 minute to pellet potassium borohydride, and decylubiquinol was transferred to a fresh tube. Assays were setup in 96-well format with each well containing 100 *μ*M of chemical dissolved in 180 *μ*L of complex III assay buffer (50 mM Tris-HCl (pH 7.5), 4 mM NaN3, 50 μM decylubiquinol, 50 μM oxidized CytC, 0.01% BSA, 0.05% Tween-20). DMSO, ethanol and antimycin A (100 mM) controls were included in parallel. The reaction was initiated by pipetting 20 μL of mitochondria suspension into each well and briefly mixing. Absorbance was measured at 550 nm in 30 s intervals over the course of 15 minutes. Total complex III enzymatic activity was determined by plotting absorbance versus time for each well and calculating the slope of the line during the linear phase of the initial rate of reaction (minutes 1-4). Antimycin-sensitive activity was calculated by subtracting the complex III activity of the antimycin control wells. Percent complex III activity for each chemical was then calculated by dividing the antimycin-sensitive activity of chemical-treated wells by that of the DMSO control wells.

Complex IV (Cytochrome C Oxidase) activity was determined by following the oxidation of CytC in the protocol outlined by Janssen and Boyle, 20 19^55^. Reduced CytC was prepared fresh by adding 1 μL increments of 0.1 M dithiothreitol to a 1 mM solution of CytC and vortexing. The solution changes colour from brown to orange/pink when CytC has been reduced. Reduction of CytC was checked by diluting a sample 50-fold and measuring the ratio of absorbance 550/560 nm (ratio > 6 indicates reduction). To setup the assay, 180 μL of complex IV buffer (50 mM KPi buffer, 60 μM reduced CytC) containing a dissolved hit compound at 100 μM was added to each well of a 96-well plate. DMSO and KCN (300 μM) controls were included in parallel. The reaction was initiated by pipetting 20 μL of mitochondria suspension into each well and briefly mixing. Absorbance was measured at 550 nm in 30 s intervals over the course of 15 minutes. Complex IV enzymatic activity was determined by plotting absorbance versus time for each well and calculating the slope of the line during the linear phase of the initial rate of reaction (minutes 1-3). Percent complex IV activity for each chemical was calculated by dividing the activity of chemical-treated wells by that of the DMSO controls.

For preliminary screens of hit compounds, data is the average of four biological replicates, each with two technical replicates. The complex I and II activity assays were also used for all follow-up dose-response experiments presented in this work. Assays were performed to cover range of final concentrations of chemical from 0.1 nM to 75 μM. For dose-responses of benzimidazole analogs, data represents the average of two biological replicates, each with two technical replicates. For all other dose-responses, data represents the average of four biological replicates, each with two technical replicates.

### *C. elegans in vivo* ATP levels

*In vivo* energy levels of *C. elegans* were estimated using a bioluminescent strain (PE255) constitutively and ubiquitously expressing firefly luciferase as previously reported (Lagido *et al.*, 2008; Luz *et al.*, 2016)^56,57^. 30 μL of M9 buffer was distributed to each well of a black flat-bottomed 96-well culture plate, and 0.5 *μ*L of chemicals or DMSO (1% v/v) controls were added to the wells with a multi-channel pipette. L1 synchronized worms obtained from an embryo preparation were plated on NGM agar seeded with OP50 *E. coli* and allowed to grow for 12 hours at 20 °C. After allowing time for foraging, worms were washed off of plates, rinsed twice in M9 buffer and diluted to a concentration of ~10 worms/μL. 20 μL of worm suspension containing ~200 animals was then added to the microplate wells. Plates were sealed with an aluminum foil seal and transferred to an incubator for 6 hours at 20 °C with shaking at 165 rpm. Following incubation, 100 μL of luminescence buffer (10 mM Na_2_PO_4_, 5 mM Citric acid, 0.5% DMSO, 0.025% Triton X-100, 50 μM Luciferin) was added to each well of the microplate and mixed via 180 rpm shaking for 2.5 minutes. Luminescence was then measured on a CLARIOstar (BMG Labtech) plate reader. Percent ATP for each dose of chemical was calculated by dividing the background corrected luminescence by that of the corresponding DMSO controls. Data represents the average of four biological replicates, each with four technical replicates.

---

**Supplemental Figure 1.**
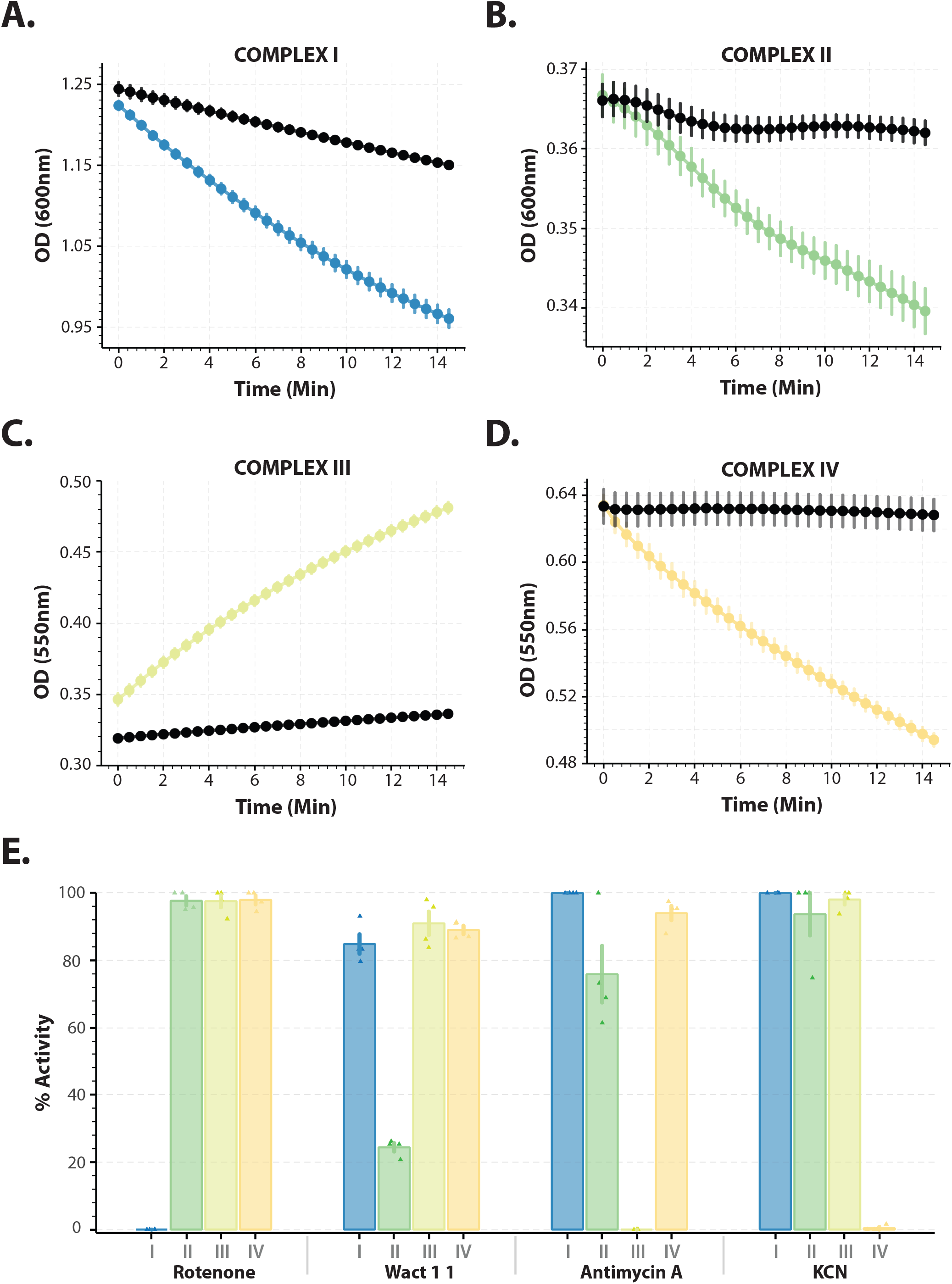
In vitro assays for each complex of the ETC. Activity assays were performed for each of the 4 core complexes of the electron transport chain using mitochondria isolated from wild-type (N2) *C. elegans:* (A) Complex I, (B) Complex II, (C) Complex III, (D) Complex IV. Activity was determined spectrophotometrically by following either the change in the reduction or oxidation of a colorimetric dye (DCIP [600nm] – Complexes I & II, Cytochrome C [550nm] – Complexes III & IV) over time; details in Materials and Methods. The initial rate of each colourimetric reaction was determined both in the presence of DMSO and a saturating dose of inhibitor to determine the specific activity of each complex in each of the respective *in vitro* assays. Percent inhibition of a given compound can thus be determined by measuring the rate of change in absorbance in mitochondria incubated with drug compared to the change in absorbance with only solvent. Data are mean of 4 independent replicates; error bars show the standard error of the mean. **E.** Known inhibitors of each of the four complexes (CI: Rotenone, CII: Wact-11, CIII: Antimycin A, CIV: KCN) were screened at 100μM in *in vitro* colourimetric assays. Percent activity of each of the 5 ETC complexes is shown for the 4 different inhibitors. Data are mean of 4 independent replicates; error bars show the standard error of the mean.

**Supplemental Figure 2:**
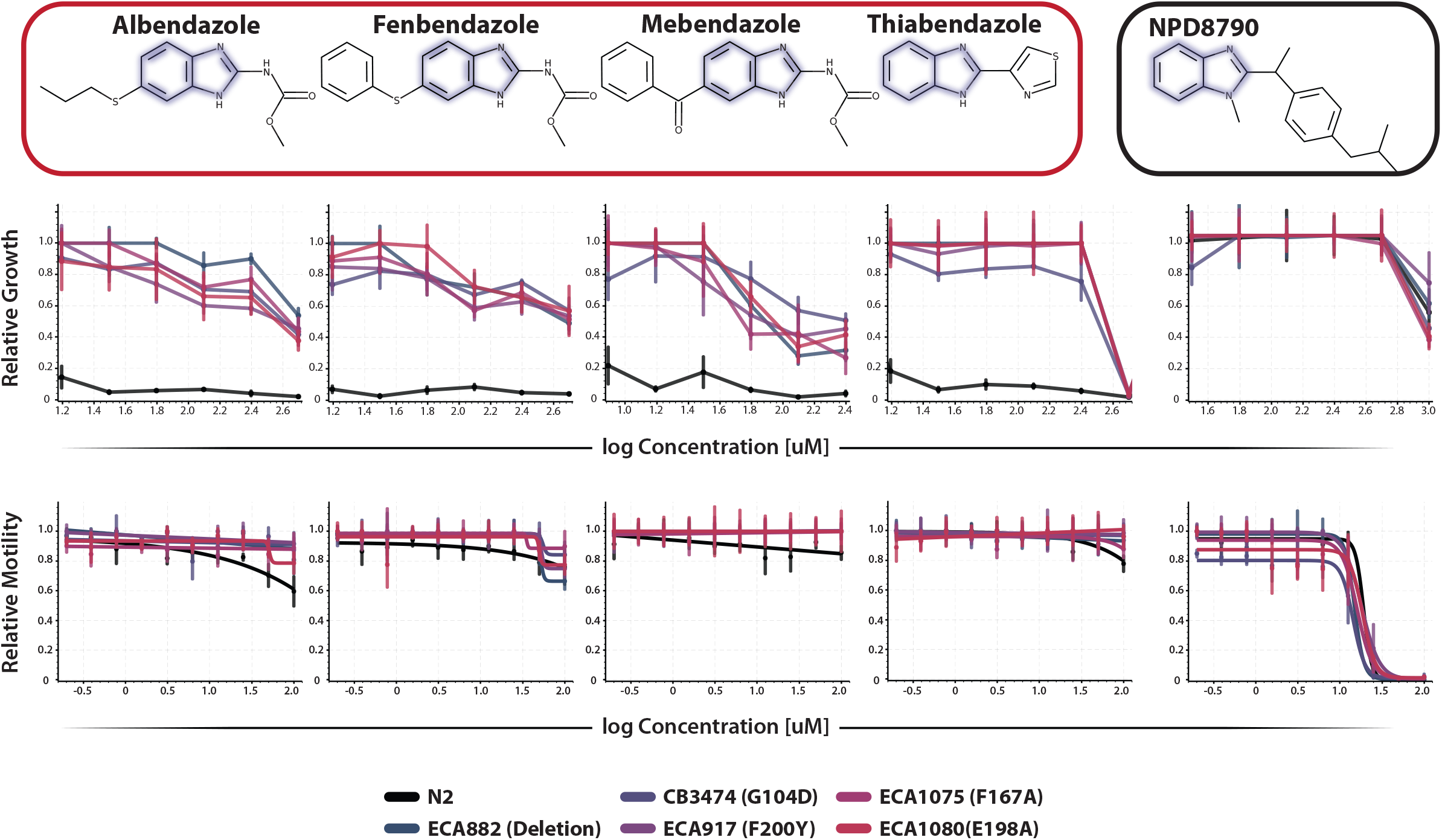
Loss of function mutations in the *C. elegans ben-1* gene confer a high degree of resistance to existing benzimidazole anthelmintics, but not to NPD8790. Growth and development dose-response curves for NPD8790 and several existing benzimidazole anthelmintics (Albendazole, Fenbendazole, Mebendazole, Thiabendazole) against wild-type (N2) and several benzimidazole-resistant (*ben-1* loss of function) worm strains. Relative growth was scored after 6-days in the presence of drug. Data are mean of 3 independent replicates; error bars show the standard error of the mean. While *ben-1* loss of function mutations allow population growth in the presence of 4 different benzimidazole anthelmintics, they do not confer any advantage to survival in the presence of NPD8790. **(Lower)** KCN survival dose-response curves for NPD8790 and several existing benzimidazole anthelmintics against wild-type (N2) and benzimidazole-resistant (*ben-1* loss of function) worm strains. Relative motility was scored 3 hours following KCN-dilution. In contrast to existing benzimidazole anthelmintics, NPD8790 elicits a strong phenotype in the KCN survival assay that is independent of *ben-1* loss of function mutations.

**Supplemental Figure 3:**
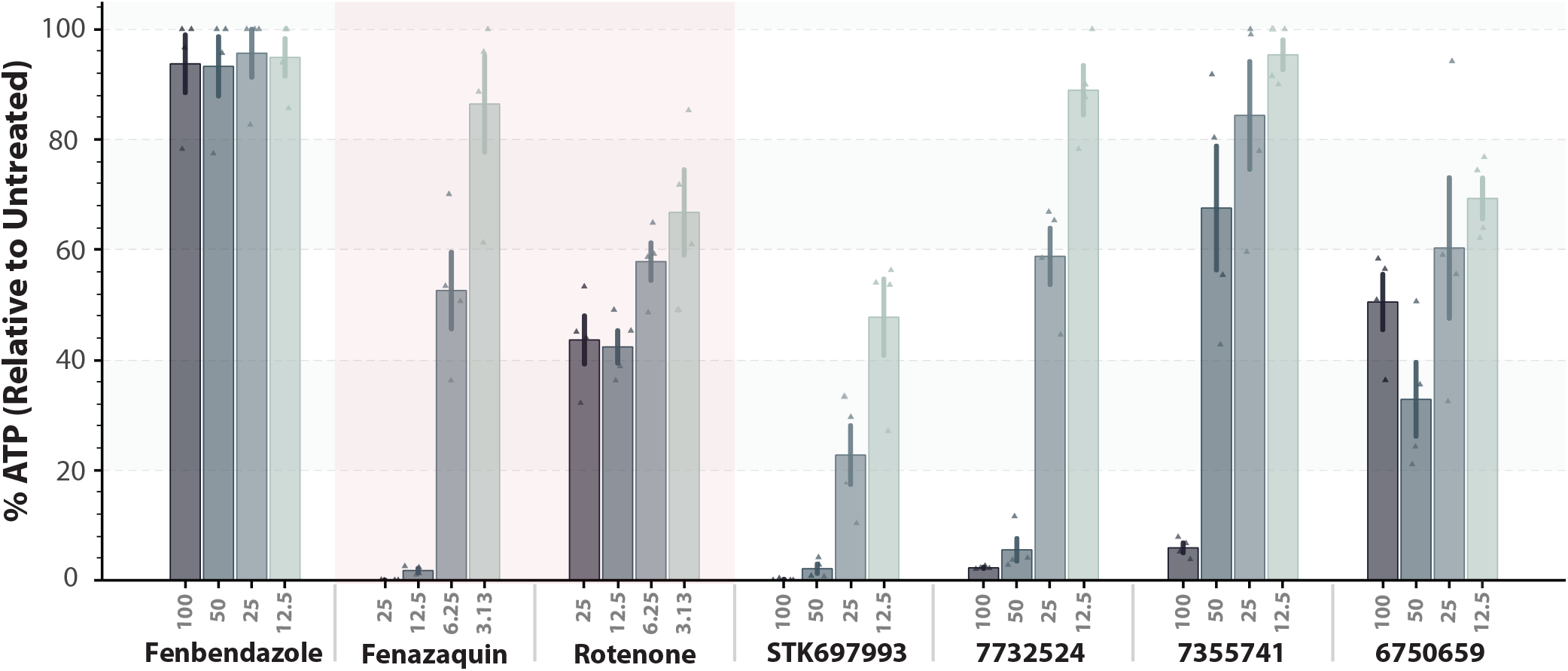
Benzimidazole analogs elicit a rapid, dose-dependent decrease in ATP levels in *C. elegans*. ATP levels of *C. elegans* L1 worms constitutively expressing a firefly luciferase construct (*Photinus pyralis)* were determined after incubation in various doses of drug for 6 hours. Percent ATP was calculated relative to untreated controls based on measured bioluminescence following the addition of D-luciferin to worms. Data are mean of 4 independent replicates; error bars show the standard error of the mean; known complex I inhibitors are displayed in red box for comparison. Similarly, to treatment with well established complex I inhibitors, incubation of *C. elegans* with benzimidazole analogs for 6 hours results in severely diminished ATP levels relative to untreated controls. In contrast, treatment with the anthelmintic benzimidazole (known to target microtubules and not the ETC) does not result in decreased ATP levels.

**Supplementary Table 1.** Bioactivity of hits identified in Riken NPDepo screens for molecules that disrupt *C. elegans* survival in anaerobic conditions (KCN).

| Cluster | Compound | Bioactivity |  |  |
| --- | --- | --- | --- | --- |
|  |  | *C. elegans* |  | HEK293 Viability |
|  |  | KCN Survival<br>EC50 (μM) | Wild type Growth<br>LC50 (μM) | LC50 (μM) |
| Control | Rotenone | 0.5 | 3.3 | ~0.79 |
| Control | Wact-11 | 2 | 0.7 | >50 |
| Authentic | Siccanin | 8 | 62.1 | 23.3 |
| Authentic | Niclosamide | <0.39 | >100 | --- |
| Authentic | Flunarizine | 16.3 | >100 | 29.3 |
| 1 | NPD6621 | 49.4 | --- | --- |
| 1 | NPD8902 | 26.6 | >100 | 21.7 |
| 2 | NPD6383 | 23.8 | --- | --- |
| 2 | NPD602 | 14.4 | --- | --- |
| 2 | NPD8034 | 12.5 | >100 | 11 |
| 2 | NPD1450 | 16.3 | >100 | >50 |
| 2 | NPD6380 | 15.6 | >100 | 21.6 |
| 3 | NPD8582 | 13.8 | --- | --- |
| 3 | NPD8298 | 2.6 | >100 | >50 |
| 3 | NPD6240 | 7.7 | >100 | 21.6 |
| 3 | FSL0005 | 0.9 | 3.4 | 6.3 |
| 3 | NPD8366 | 11.9 | >100 | 18.8 |
| 4 | NPD10504 | 12.5 | >100 | 26.1 |
| 4 | NPD3577 | 12.5 | >100 | 20.9 |
| 4 | NPD6303 | 25.5 | >100 | 38.5 |
| 4 | NPD8790 | 14.5 | >100 | >50 |
| 5 | NPD10211 | 17.7 | --- | --- |
| 5 | NP974 | 21.4 | --- | --- |
| 5 | NPD390 | 17 | 48.9 | >50 |
| 6 | NPL50654-01 | 8.1 | 26.1 | 3.8 |
| 7 | 6-heptadeca-9Z,12Z-dienyl<br>salicylic acid | 3.5 | --- | --- |
| 7 | Anacardic Acid | 5.6 | >100 | >50 |
| Growth | NPD5176 | --- | 94 | >50 |
| Growth | NPD5219 | --- | 43.02 | 16.7 |
| Growth | HTD0465 | --- | 38.2 | 18.8 |
| Growth | STK418118 | --- | 44.1 | >50 |

## REFERENCES

1. Torgerson, P. R. et al. World Health Organization Estimates of the Global and Regional Disease Burden of 11 Foodborne Parasitic Diseases, 2010: A Data Synthesis. PLoS Med 12, e1001920 (2015).

2. Pullan, R. L., Smith, J. L., Jasrasaria, R. & Brooker, S. J. Global numbers of infection and disease burden of soil transmitted helminth infections in 2010. Parasit. Vectors 7, 37 (2014).

3. Geerts, S. & Gryseels, B. Anthelmintic resistance in human helminths: a review. Trop. Med. Int. Health 6, 915–921 (2001).

4. Berry, B. J., Baldzizhar, A., Nieves, T. O. & Wojtovich, A. P. Neuronal AMPK coordinates mitochondrial energy sensing and hypoxia resistance in C. elegans. FASEB J. 34, 16333–16347 (2020).

5. Wolstenholme, A. J., Fairweather, I., Prichard, R., Von Samson-Himmelstjerna, G. & Sangster, N. C. Drug resistance in veterinary helminths. Trends Parasitol. 20, 469–476 (2004).

6. Tielens, A. G. M. Energy generation in parasitic helminths. Parasitol. Today 10, 346–352 (1994).

7. Tielens, A. G. M. & Van Hellemond, J. J. The electron transport chain in anaerobically functioning eukaryotes. Biochim. Biophys. Acta - Bioenerg. 1365, 71–78 (1998).

8. Kita, K. Electron-transfer complexes of mitochondria in Ascaris suum. Parasitol. Today 8, 155–159 (1992).

9. Kita, K., Nihei, C. & Tomitsuka, E. Parasite Mitochondria as Drug Target: Diversity and Dynamic Changes During the Life Cycle. Curr. Med. Chem. 10, 2535–2548 (2003).

10. Allen, P. C. Helminths: Comparison of their rhodoquinone. Exp. Parasitol. 34, 211–219 (1973).

11. Del Borrello, S. et al. Rhodoquinone biosynthesis in C. elegans requires precursors generated by the kynurenine pathway. eLife 8, 1–21 (2019).

12. Osada, H. & Nogawa, T. Systematic isolation of microbial metabolites for natural products depository (NPDepo). Pure Appl. Chem. 84, 1407–1420 (2011).

13. Kato, N., Takahashi, S., Nogawa, T., Saito, T. & Osada, H. Construction of a microbial natural product library for chemical biology studies. Curr. Opin. Chem. Biol. 16, 101–108 (2012).

14. Driscoll, M., Dean, E., Reilly, E., Bergholz, E. & Chalfie, M. Genetic and molecular analysis of a Caenorhabditis elegans beta-tubulin that conveys benzimidazole sensitivity. J Cell Biol 109, 2993–3003 (1989).

15. Enos, A. & Coles, G. C. Effect of benzimidazole drugs on tubulin in benzimidazole resistant and susceptible strains of Caenorhabditis elegans. Int. J. Parasitol. 20, 161–167 (1990).

16. Ghisi, M., Kaminsky, R. & Mäser, P. Phenotyping and genotyping of Haemonchus contortus isolates reveals a new putative candidate mutation for benzimidazole resistance in nematodes. Vet. Parasitol. 144, 313–320 (2007).

17. Hahnel, S. R. et al. Extreme allelic heterogeneity at a Caenorhabditis elegans beta-tubulin locus explains natural resistance to benzimidazoles. PLOS Pathog. 14, e1007226 (2018).

18. World Health Organization. Prevention and control of schistosomiasis and soil-transmitted helminthiasis: report of a WHO expert committee. (2002).

19. Lacey, E. Mode of action of benzimidazoles. Parasitol. Today 6, 112–115 (1990).

20. Brown, H. D. et al. ANTIPARASITIC DRUGS. IV. 2-(4’-THIAZOLYL)-BENZIMIDAZOLE, A NEW ANTHELMINTIC. J. Am. Chem. Soc. 83, 1764–1765 (1961).

21. Campbell, W. C., Fisher, M. H., Stapley, E. O., Albers-Schönberg, G. & Jacob, T. A. Ivermectin: a potent new antiparasitic agent. Science 221, 823–828 (1983).

22. Campbell, W. C. History of avermectin and ivermectin, with notes on the history of other macrocyclic lactone antiparasitic agents. Curr. Pharm. Biotechnol. 13, 853–865 (2012).

23. Kita, K. [Current Trend of Drug Development for Neglected Tropical Diseases (NTDs)]. Yakugaku Zasshi 136, 205–11 (2016).

24. Moore, H. W. & Folkers, K. Coenzyme Q. LXII. Structure and Synthesis of Rhodoquinone, a Natural Aminoquinone of the Coenzyme Q Group. J. Am. Chem. Soc. 87, 1409–1410 (1965).

25. Ozawa, H., Sato, M., Natori, S. & Ogawa, H. Rhodoquinone-9 from the Muscle of Ascaris lumbricoides var. suis. Chem. Pharm. Bull. (Tokyo) 18, 1099–1103 (1970).

26. Sato, M. & Ozawa, H. Occurrence of ubiquinone and rhodoquinone in parasitic nematodes, metastrongylus elongatus and Ascaris lumbricoides var. suis. J. Biochem. (Tokyo) 65, 861–867 (1969).

27. Tan, J. H. et al. Alternative splicing of coq-2 controls the levels of rhodoquinone in animals. eLife 9, e56376 (2020).

28. Roberts Buceta, P. M. et al. The kynurenine pathway is essential for rhodoquinone biosynthesis in Caenorhabditis elegans. J. Biol. Chem. 294, 11047–11053 (2019).

29. Blaxter, M. Caenorhabditis elegans is a nematode. Science 282, 2041–2046 (1998).

30. Hoebeke, J., Van Nijen, G. & De Brabander, M. Interaction of oncodazole (R 17934), a new antitumoral drug, with rat brain tubulin. Biochem. Biophys. Res. Commun. 69, 319–324 (1976).

31. Tamaoki, T. et al. Staurosporine, a potent inhibitor of phospholipid/Ca++dependent protein kinase. Biochem. Biophys. Res. Commun. 135, 397–402 (1986).

32. Hsiang, Y. H., Hertzberg, R., Hecht, S. & Liu, L. F. Camptothecin induces protein-linked DNA breaks via mammalian DNA topoisomerase I. J. Biol. Chem. 260, 14873–14878 (1985).

33. Chen, B. et al. Inhibitory mechanism of reveromycin A at the tRNA binding site of a class I synthetase. Nat. Commun. 12, 1616 (2021).

34. Anger, E. E., Yu, F. & Li, J. Aristolochic Acid-Induced Nephrotoxicity: Molecular Mechanisms and Potential Protective Approaches. Int. J. Mol. Sci. 21, E1157 (2020).

35. Pearson, R. D. & Hewlett, E. L. Niclosamide therapy for tapeworm infections. Ann. Intern. Med. 102, 550–551 (1985).

36. Frayha, G. J., Smyth, J. D., Gobert, J. G. & Savel, J. The mechanisms of action of antiprotozoal and anthelmintic drugs in man. Gen. Pharmacol. 28, 273–299 (1997).

37. Mogi, T. et al. Siccanin rediscovered as a species-selective succinate dehydrogenase inhibitor. J. Biochem. (Tokyo) 146, 383–387 (2009).

38. Nose, K. & Endo, A. Mode of action of the antibiotic siccanin on intact cells and mitochondria of Trichophyton mentagrophytes. J. Bacteriol. 105, 176–184 (1971).

39. Burns, A. R. et al. Caenorhabditis elegans is a useful model for anthelmintic discovery. Nat Commun 6, 7485 (2015).

40. Burns, A. R. et al. High-throughput screening of small molecules for bioactivity and target identification in Caenorhabditis elegans. Nat Protoc 1, 1906–14 (2006).

41. Triner, L., Vulliemoz, Y., Schwartz, I. & Nahas, G. G. Cyclic phosphodiesterase activity and the action of papaverine. Biochem. Biophys. Res. Commun. 40, 64–69 (1970).

42. Benej, M. et al. Papaverine and its derivatives radiosensitize solid tumors by inhibiting mitochondrial metabolism. Proc. Natl. Acad. Sci. U. S. A. 115, 10756–10761 (2018).

43. Morikawa, N., Nakagawa-Hattori, Y. & Mizuno, Y. Effect of dopamine, dimethoxyphenylethylamine, papaverine, and related compounds on mitochondrial respiration and complex I activity. J. Neurochem. 66, 1174–1181 (1996).

44. Dilks, C. M. et al. Quantitative benzimidazole resistance and fitness effects of parasitic nematode beta-tubulin alleles. Int. J. Parasitol. Drugs Drug Resist. 14, 28–36 (2020).

45. Dilks, C. M., Koury, E. J., Buchanan, C. M. & Andersen, E. C. Newly identified parasitic nematode beta-tubulin alleles confer resistance to benzimidazoles. Int. J. Parasitol. Drugs Drug Resist. 17, 168–175 (2021).

46. Lautens, M. J. et al. Identification of enzymes that have helminth-specific active sites and are required for Rhodoquinone-dependent metabolism as targets for new anthelmintics. PLoS Negl. Trop. Dis. 15, e0009991 (2021).

47. Nakagawa, Y., Kuwano, E., Eto, M. & Fujita, T. Effects of Insect-Growth-Regulatory Benzimidazole Derivatives on Cultured Integument of the Rice Stem Borer and Mitochondria from Rat Liver. Agric. Biol. Chem. 49, 3569–3573 (1985).

48. Stiernagle, T. Maintenance of C. elegans. WormBook 1–11 (2006) doi:10.1895/wormbook.1.101.1.

49. Spensley, M., Del Borrello, S., Pajkic, D. & Fraser, A. G. Acute Effects of Drugs on Caenorhabditis elegans Movement Reveal Complex Responses and Plasticity. G3 Bethesda Md 8, 2941–2952 (2018).

50. Sander, T., Freyss, J., von Korff, M. & Rufener, C. DataWarrior: an open-source program for chemistry aware data visualization and analysis. J. Chem. Inf. Model. 55, 460–473 (2015).

51. O’Boyle, N. M., Morley, C. & Hutchison, G. R. Pybel: a Python wrapper for the OpenBabel cheminformatics toolkit. Chem. Cent. J. 2, 5 (2008).

52. Cline, M. S. et al. Integration of biological networks and gene expression data using Cytoscape. Nat. Protoc. 2, 2366–2382 (2007).

53. Long, J. et al. Comparison of two methods for assaying complex I activity in mitochondria isolated from rat liver, brain and heart. Life Sci. 85, 276–280 (2009).

54. Luo, C., Long, J. & Liu, J. An improved spectrophotometric method for a more specific and accurate assay of mitochondrial complex III activity. Clin. Chim. Acta Int. J. Clin. Chem. 395, 38–41 (2008).

55. Janssen, R. C. & Boyle, K. E. Microplate Assays for Spectrophotometric Measurement of Mitochondrial Enzyme Activity. Methods Mol. Biol. Clifton NJ 1978, 355–368 (2019).

56. Lagido, C., Pettitt, J., Flett, A. & Glover, L. A. Bridging the phenotypic gap: real-time assessment of mitochondrial function and metabolism of the nematode Caenorhabditis elegans. BMC Physiol. 8, 7 (2008).

57. Luz, A. L., Lagido, C., Hirschey, M. D. & Meyer, J. N. In Vivo Determination of Mitochondrial Function Using Luciferase-Expressing Caenorhabditis elegans: Contribution of Oxidative Phosphorylation, Glycolysis, and Fatty Acid Oxidation to Toxicant-Induced Dysfunction. Curr. Protoc. Toxicol. 69, 25.8.1–25.8.22 (2016).

